# Empowering multiplexed ultra-throughout ribosome profiling with RiboWich

**DOI:** 10.1101/2025.10.17.683053

**Authors:** Ilaria Bruno, Cecilia Perrucci, Gabriele Tomè, Giorgia Susin, Stefania Mazzalai, Martina Sevegnani, Alessia Del Piano, Luisa Donini, Manuela Basso, Massimiliano Clamer, Elena Perenthaler, Fabio Lauria, Gabriella Viero

## Abstract

Ribosome profiling (RiboSeq) improved the understanding of mRNA translation, enabling the precise mapping of ribosome positioning along transcripts at single-nucleotide resolution. Although various library preparation protocols overcome the need for high input material, their technical complexity and limited efficiency hinder robust profiling, preventing them from keeping pace with other sequencing techniques and applications. To move towards high-throughput and single-cell RiboSeq technologies, we developed RiboWich (<u>Ribo</u>some sand<u>Wich</u>). By directly ligating adaptors to ribosome-embedded RNA fragments, RiboWich eliminates the need for ribosome purification and size-selecting ribosome footprints and tackles two major bottlenecks, expanding RiboSeq for more advanced technologies. RiboWich offers robustness in profiling and excels in detecting upstream translons in immortalized and primary cells alike. By exploiting a dual-step multiplexing strategy, RiboWich enables the simultaneous profiling of at least 96 samples while retaining sensitivity and capturing condition-specific differences in translation. By enabling scalable, low-input translatome profiling, this advancement empowers proteogenomic approaches and AI/ML-driven data analysis to uncover regulatory dynamics, neoantigens, functional small translons, and drug-responsive signatures across diverse biological contexts and users. Altogether, RiboWich represents a straightforward, versatile, and scalable ribosome profiling conceptual platform that combines accessibility, sensitivity, and throughput potential, laying the foundation for advanced single-cell RiboSeq applications.

## Introduction

Several experimental approaches investigate the complete set of mRNAs engaged in translation, in both cell lines and tissues (King & Gerber, 2016; Ingolia, 2019; Zhao et al., 2019). Expanding on the pioneering ribosome footprinting approach by Wolin and Walter able to generate RNA fragments shielded by ribosomes from endonuclease digestion (Wolin and Walter, 1988), Ingolia and colleagues proposed a seminal library preparation protocol to couple ribosome footprinting with next-generation sequencing: ribosome profiling (RiboSeq) (Ingolia et al., 2009). Profiling the landscape of ribosomes on RNA at single-nucleotide resolution has proved to be of paramount relevance, as ribosome localization may reveal uncharacterized layers of gene expression regulation. These insights expand our knowledge of conventional translons (TLs), *i.e*., Open Reading Frames (ORFs) showing evidence of translation (Goel et al., 1973; Tierney et al., 2024; Świrski et al., 2025), as well as non-conventional TLs, including upstream and downstream TLs and, more generally, of the peptide-coding potential of the transcriptome (Calviello et al., 2016; Ingolia, 2016; Brar & Weissman, 2015; Chothani et al., 2022).

RiboSeq is based on the generation of around 30 nt-long Ribosome Protected Fragments (RPFs) followed by their isolation and conversion into a dsDNA library suitable for sequencing (Ingolia et al., 2009; Ingolia et al., 2012). RiboSeq generates information-rich data that can be used for diverse purposes, including: i) map the position of ribosomes along transcripts with single-nucleotide resolution (Lauria et al., 2018; Tierney et a., 2024), ii) provide quantitative estimations of the engagement of ribosomes on mRNAs (Ingolia et al., 2012; Ingolia, 2016), iii) identify RNAs with canonical or non-canonical coding potential (Calviello et al., 2016, Ingolia, 2016; Brar & Weissman, 2015; Chothani et al., 2022, Ingolia et al., 2011), iv) determine the levels of potential ribosome-based filtering control of mRNA translation (Mauro and Edelman, 2002) when coupled with immunoprecipitation of specific ribosomal proteins (Simsek et al., 2017, Susanto et al., 2024), ribosome-associated proteins (Lauria et al., 2020) or cell type-specific ribosomal populations (Heiman et al., 2008; Doyle & Dougherty et al., 2008), v) identify the population of mRNAs undergoing active translation (Clamer et al., 2018), vi) study translation fidelity (Wang and Mao, 2019), vi) determine gene-specific translation elongation rates (Ingolia et al., 2011; Gerashchenko et al., 2021), and others.

Although widely applied, RiboSeq presents substantial challenges, often reflected in the sub-optimal data quality (Wang and Mao, 2023). It remains a complex, labor-intensive, and low-throughput technique that typically requires large quantities of starting material to yield sufficient RPFs for successful library preparation (Wang and Mao, 2023; Tumuro and Iwasaki, 2025). Over the years, various protocol steps have been refined, including cell lysis (McGlincy & Ingolia, 2017), RNase digestion (Gerashchenko and Gladyshev, 2017), ribosome isolation (Gerashchenko and Gladyshev, 2017; Zhang et al., 2017; Clamer et al., 2018), and, particularly, library construction. For example, circularization-based methods involve RNA ligation and DNA circularization and are hindered by inefficiency and 5’ end inaccuracies (Ingolia et al., 2009; McGlincy & Ingolia, 2017). Dual-ligation methods have been proposed to address these issues, requiring ligation of 3’ and 5’ adaptors to RNA fragments (VanIsberghe et al., 2021). However, challenges like ligation biases and adaptor self-ligation strongly reduce the overall efficiency. More recently, template switch-based methods have gained traction owing to their ability to handle low-input samples (Li et al. 2022, 2023; Xiong et al. 2022; Zhang et al. 2022; Zou et al. 2022). Despite their potential, these approaches introduce variability and strong sequence biases at the 5’ end of each RPF, which impact the accuracy of ribosome occupancy assessment. While both dual-ligation and template-switching methods are undoubtedly promising for low-input (Hornstein, N. et al., 2016; Xiong et al., 2022; Li et al., 2022; Zhang et al.,2022; Meindl et al., 2023; Froberg et al.,2023; Ozadam et al., 2023) and single-cell (VanIsberghe et al., 2021) applications, addressing biases and optimizing each step remains crucial for generating reliable, high-quality datasets. Altogether, overcoming these issues with novel methods is key to advancing translation analysis into high-throughput research and clinical settings, thereby fueling data demands of next-generation Artificial Intelligence/Machine Learning technologies.

Here, we introduce RiboWich (<u>Ribo</u>some sand<u>Wich</u>), an innovative protocol that reshapes RiboSeq by addressing previously overlooked steps: ribosome isolation, RNA extraction, and, when required by the protocol, also RPF size selection. At the core of RiboWich lies a direct, in-lysate ligation of sequencing adaptors to the Ribosome-Embedded RNA Fragments (REFs), completely bypassing the need for ribosome isolation, RNA extraction, and RPF size selection. RiboWich represents a streamlined, versatile, and scalable ribosome profiling approach enabling applications ranging from challenging, low-input samples to high-throughput experiments.

## Results

### RiboWich captures bona fide RPFs in HEK293T

A key step in RiboSeq experiments is the generation and isolation of Ribosome Protected Fragments (RPFs) (**Figure 1A**, left). RPFs are generated by treating cytoplasmic lysates with an endonuclease, typically RNase I, which digests unprotected RNA, leaving ribosome footprints with 5’-hydroxyl (5’-OH) and 3’-phosphate (3’-P) ends, without positional cleavage bias (Ingolia et al., 2009). Upon nuclease treatment, protocols proceed with ribosome isolation, RNA extraction, and size selection of the RPFs through polyacrylamide gel electrophoresis (Ingolia et al., 2009). Next, the purified ∼30 nt-long ‘naked’ fragments, bearing a 3’-P group, are used to prepare a sequencing library through a series of protocol-specific enzymatic reactions, typically starting with a 3’-dephosphorylation. Finally, the library is amplified and sequenced (Wang and Mao, 2023). Ribosome profiling protocols remain lengthy and technically demanding, and simplifying the workflow is key to fostering the development of rapid, automated protocols for high-throughput and high-resolution ribosome profiling. Leveraging the enzymatic capability of RtcB-like enzymes to directly ligate RNA fragments with a 3’-P (Popow et al., 2011; Moncan et al., 2023) to convenient adaptors (Del Piano et al., 2021), we developed RiboWich, a streamlined solution that enables a direct, in-lysate adaptor ligation to RNA fragments while they are still embedded within ribosomes (**Figure 1A**, right). Upon adaptor ligation, the remaining steps to generate a dsDNA library are performed according to a commercial kit (see Methods) for sequencing on the Illumina platform. By performing the adaptor ligation as the very first reaction, bypassing ribosome isolation, RNA extraction, RPF isolation, and 3’ dephosphorylation, RiboWich easily integrates with multiplexing strategies. To fully exploit this advantage, we implemented a dual-step multiplexing approach that enables the simultaneous processing of 96 samples (**Figure 1B**). Sample preparation is carried out in plate, beginning with lysis and RNase I digestion. Next, in the first barcoding step, barcoded adaptors are ligated directly to REFs, allowing 96 wells to be pooled into 12 tubes. After 5′ phosphorylation and circularization, a second barcoding step is performed using barcoded reverse transcription (RT) primers, uniquely labeling each sample and thereby allowing all libraries to be combined into a single tube for PCR amplification. This streamlined workflow minimizes handling, enabling scalable, high-throughput ribosome profiling.

**Figure 1.**
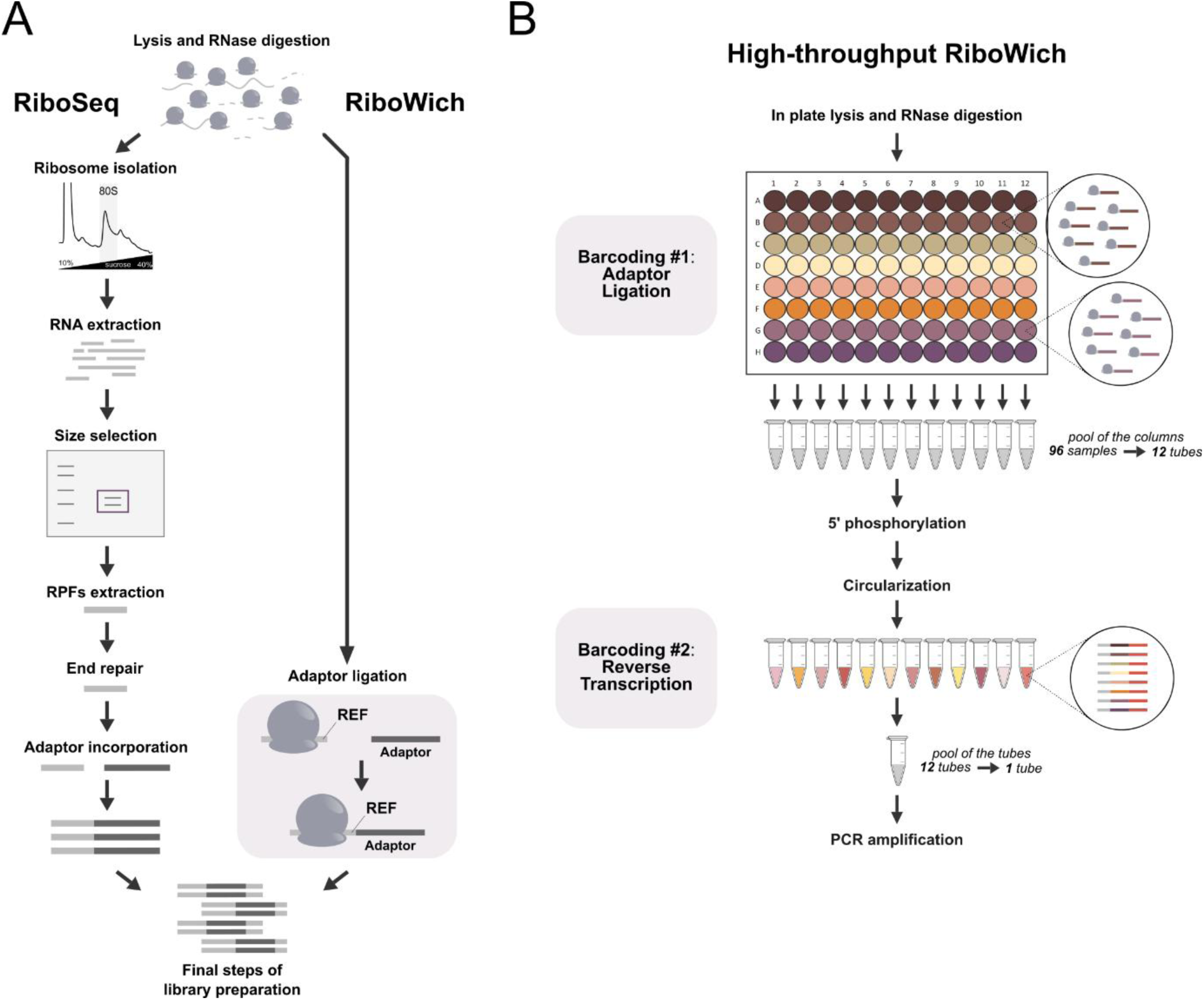
RiboWich and high-throughput RiboWich for ribosome profiling. (**A**) Schematic comparison of conventional RiboSeq (left) and RiboWich (right) protocols. In both approaches, cytoplasmic lysates are treated with RNase to digest free RNA not protected by ribosomes. All conventional RiboSeq protocols then require ribosome isolation, RNA extraction, polyacrylamide gel size selection, and recovery of RPFs. With RiboWich, the adaptor is directly ligated to the ribosome-embedded RNA fragments (REFs) within the lysate, skipping the abovementioned steps. Finally, both protocols proceed with the following steps of library preparation. (**B**) Schematic of high-throughput RiboWich with a dual-step multiplexing strategy. Lysis and RNase digestion are performed in plate. Ligation serves as the first barcoding step, where barcoded adaptors are ligated to REFs, enabling pooling of 96 individual wells into 12 tubes. After 5’ phosphorylation and circularization, the second step of barcoding is carried out: barcoded reverse transcription (RT) primers are exploited, allowing all samples to be combined into a single tube for PCR amplification.

First, we demonstrated the effectiveness of the *in vitro* adaptor ligation to the REFs and showed that the RPF purification is not essential for ribosome profiling. To this aim, we compared RiboWich to a standard method (hereafter referred to as RiboSeq) (**Supplementary Figure 1A**), which includes gel-based size selection of RPFs and 5’ phosphorylation of RNA fragments prior to adaptor ligation. First, we isolated ribosomes from HEK293T cells upon RNase I treatment (**Supplementary Figure 1B**). For RiboSeq, we extracted RNA, size-selected RPFs (∼30 nt) through denaturing gel electrophoresis (**Supplementary Figure 1C**), and prepared the library with the same chemistry used for RiboWich (see Methods). In parallel, we performed RiboWich (**Supplementary Figure 1D**). After precipitating and resuspending the ribosomes in an optimized *in vitro* Ribosome Solution, we ligated the adaptor to the REFs. To enable a direct comparison between the two workflows, we introduced a gel-based purification step of the ligation products (**Supplementary Figure 1E**). Next, we proceeded with the library preparation using a commercial kit (see Methods). In both cases, we obtained a library of the expected size (∼200 bp) after PCR amplification, suggesting successful adaptor ligation to the REFs (**Supplementary Figure 1F-G**).

After sequencing, we performed a comprehensive quantitative assessment of both RiboSeq and RiboWich datasets, analyzing various performance indicators as quality measures (Lauria et al., 2018). First, the proportion of reads aligning to the transcriptome is higher in RiboWich (mean = 16.5%) compared to RiboSeq (mean = 12.6%) (**Figure 2A** and **Supplementary Table 1**). The mean percentage distribution of read lengths peaks at the expected RPF dimension (30-34 nt) in both methods (**Figure 2B**). RiboWich shows a slightly broader range, extending into the 20-26 nt region, which may represent diverse ribosome conformations (Lareau et al., 2014; Wang and Mao, 2023; Wu et al., 2019). RPFs are expected to be predominantly located within the coding sequence (CDS) of mRNAs, where the ribosome P-sites, taken as a reference position, are expected to exhibit trinucleotide periodicity. To assess the enrichment of ribosome occupancy in the CDS, we analyzed the percentage of P-sites mapping on the 5’ untranslated region (UTR), CDS, and 3’ UTR portions of mRNAs (**Figure 2C**, left and middle), and compared to a uniform distribution weighted on region lengths with random P-site positioning along mRNAs (**Figure 2C**, right). In RiboSeq and RiboWich, the percentage of P-sites mapping on each mRNA region is similar, showing a strong enrichment in the CDS.

**Figure 2.**
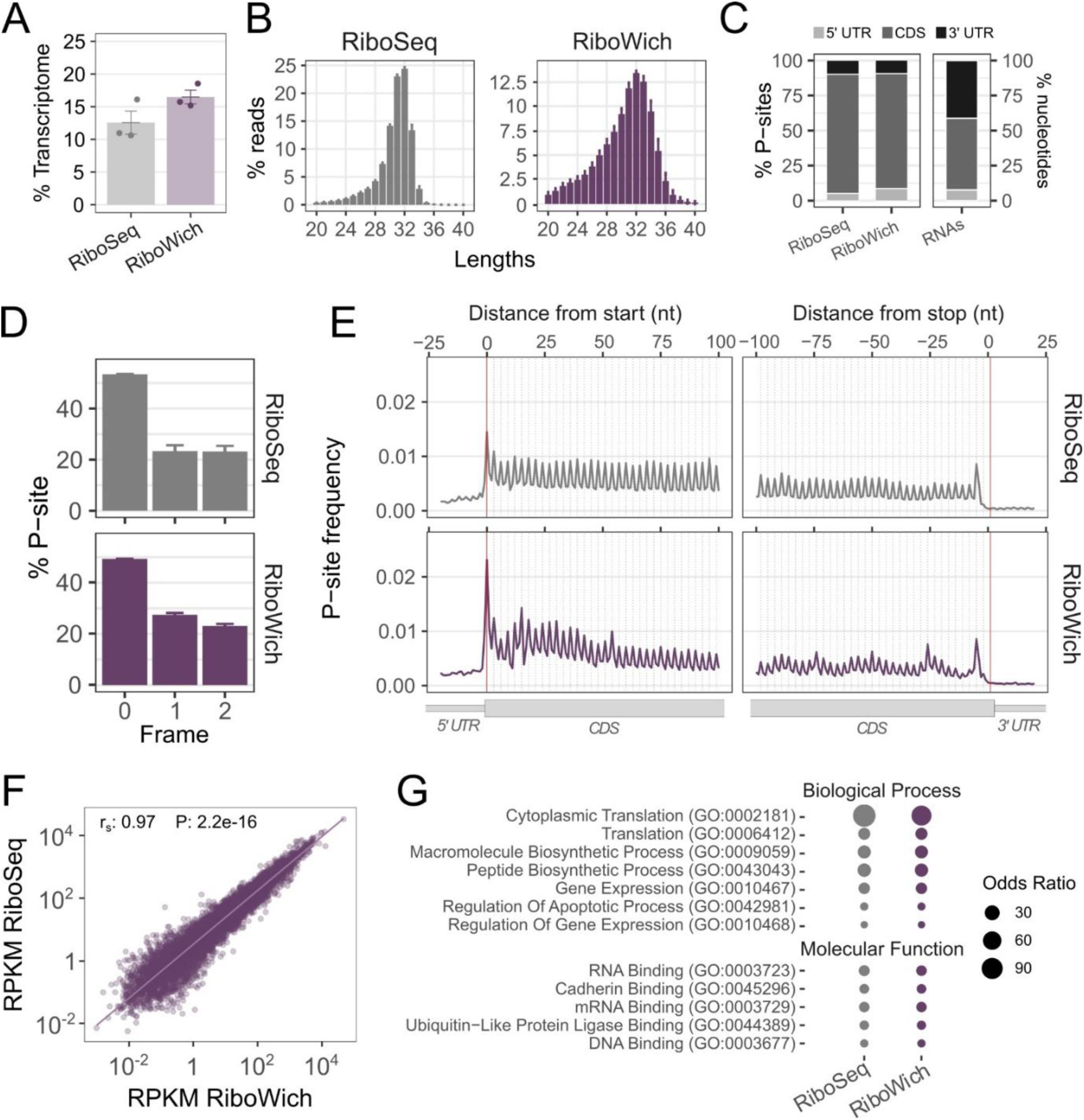
Comparison between RiboSeq and RiboWich in HEK293T. (**A**) Percentages of reads aligning to the transcriptome for RiboSeq and RiboWich. Results represent the mean ± SEM of n=3 independent technical replicates. Two-sided T-test: not significant. (**B**) Percentage distribution of read lengths for RiboSeq (left) and RiboWich (right). Results represent the mean ± SEM of n=3 independent technical replicates. (**C**) Percentages of P-sites mapping on the 5’ UTR, CDS, and 3’ UTR for RiboSeq (left) and RiboWich (right). Percentages of region lengths in mRNA sequences are reported as reference (right). Results represent the mean of n=3 independent technical replicates. (**D**) Percentage of P-sites mapping to the three reading frames for RiboSeq (top) and RiboWich (bottom). Results represent the mean ± SEM of n=3 independent technical replicates. (**E**) Meta-profiles showing the frequency of P-sites mapping at the beginning and at the end of the coding sequence in RiboSeq (top) and RiboWich (bottom). Results represent the mean ± SEM of n=3 independent technical replicates. (**F**) Correlations of RPKM per protein-coding genes (n = 20496) between RiboSeq and RiboWich. The Spearman correlation and the statistical significance based on the two-tailed William’s test are reported. (**G**) Enrichment analysis based on Gene Ontology Biological Process (top) and Molecular Function (bottom) terms of the top covered 5% genes. All terms display a statistically significant enrichment (adjusted p-value < 0.05), and the dots’ size is proportional to the Odds Ratio.

In addition, the majority of the P-sites in the CDS fall in the correct reading frame (0) to a similar extent in both methods (**Figure 2D**). Next, we visually inspected the trinucleotide periodicity of P-sites mapping along the CDS, around the start and stop codons, also including portions of the UTRs. As expected for high-quality ribosome profiling data, both RiboSeq and RiboWich show sharp peaks associated with in-frame periodicity along the CDS and low signal on the UTRs (**Figure 2E**), demonstrating the success of the ligation to the REFs. Importantly, a direct comparison of the ribosome occupancy expressed as Reads Per Kilobase of transcript per Million mapped reads (RPKM) shows a strong correlation between the methods (**Figure 2F** and **Supplementary Table 2**). To further reinforce our findings, we performed Gene Ontology enrichment analysis of mRNAs captured by the two methods and found similar results (**Figure 2G** and **Supplementary Table 3**).

Overall, these results demonstrate that *in vitro* RiboWich performs at least as efficiently as established ribosome profiling protocols and is capable of capturing *bona fide* RPFs, providing information on ribosome positioning on mRNAs across the translatome.

### RiboWich uncovers non-canonical translation events occurring in the 5’ UTR of mRNAs

To explore RiboWich’s potential and sensitivity, we investigated non-canonical translation events, focusing on translons located upstream of canonical initiation sites, *i.e.*, non-overlapping and overlapping upstream translons (uTLs). Despite being present in 40-50% of mammalian transcripts, uTLs remain largely unannotated as translationally active sites (Wright et al., 2022). Nonetheless, they are attracting growing attention for their potential clinical relevance (Dasgupta et al., 2024), including in viral translation, cancer drug resistance, and rare genetic diseases (Silva et al., 2019; Khan and Fox, 2024). Existing ribosome profiling methods have been considered not sensitive enough to capture these events with high probability and resolution (Wang and Mao, 2023), and proteomics methods often struggle to identify short (< 100 amino acids), unstable peptides derived from non-canonical events (Oyama et al., 2007; Slavoff et al., 2013). These limitations underscore the importance of high-quality RiboSeq data to accurately distinguish novel uTLs from background noise, making this an excellent test case for showcasing RiboWich’s performance compared to RiboSeq.

We applied RiboSeq and RiboWich in HEK293T treated with 2 μg/mL harringtonine for 10 minutes (harringtonine-treated, HT, **Supplementary Figure 2A-C**), a molecule that inhibits translation initiation and stalls ribosomes at translation start sites (Huang 1975), allowing both the identification of annotated coding regions and the discovery of novel, unannotated translons (Ingolia et al., 2011; Gerashchenko et al., 2021; Tong and Martinez, 2025). We observed that RiboWich shows a higher percentage of reads aligned to the transcriptome (mean = 6.02%) compared to RiboSeq (mean = 3.52%) (**Supplementary Figure 2D** and **Supplementary Table 1**). The mean percentage distribution of read lengths peaks at the expected RPF dimensions (30-34 nt) in both methods (**Supplementary Figure 2E**), and the majority of P-sites in the CDS fall in the correct reading frame (0) (**Supplementary Figure 2F**). The trinucleotide periodicity of the P-sites across the meta-profiles shows an increase (RiboSeq: 11.9% and RiboWich: 4.6%) in the P-sites frequency at the start codon and a relative decrease along the CDS in both RiboSeq and RiboWich datasets (**Supplementary Figure 2G**).

While both RiboSeq and RiboWich preserve signal along the CDS, the percentage of P-sites mapping on the 5’ UTR is generally high, implying the presence of non-canonical initiation events. RiboWich exhibits a significant enrichment of P-sites in the 5’ region compared to RiboSeq (**Figure 3A**), suggesting that RiboWich is likely more effective than RiboSeq at capturing non-canonical translation events occurring in the 5’ UTR. To dissect whether RiboWich improves the detection of these alternative events, we developed and implemented a dedicated computational pipeline, based on (**Figure 3B**). First, our workflow identifies all potential uTLs in the 5’ UTR, characterized by a start codon and a stop codon in frame with respect to the start site. Then, our pipeline applies three sequential filters to minimize false positives and accurately select only uTLs characterized by: i) more than 10 RPKM; ii) more than 25% of the reads mapping at the start codon in the HT condition, as expected due to ribosome accumulation at initiation sites; iii) trinucleotide periodicity in NT samples. By intersecting the resulting uTLs with the RibouORF database (Liu et al., 2023), we identified 502 uTLs in RiboSeq and 601 in RiboWich. Interestingly, in RiboWich, the number of uTLs, normalized on the total TLs, is significantly higher compared to RiboSeq (**Figure 3C**), highlighting RiboWich’s superior ability to capture uTLs compared to standard RiboSeq. The detected uTLs displayed robust trinucleotide periodicity in both NT and HT conditions and methods (**Figure 3D-E**). Importantly, RiboWich shows a significantly higher in-frame P-site signal compared to RiboSeq (**Figure 3D**), a more distinct trinucleotide periodicity in the meta-profiles around the translation initiation region in the NT condition (**Figure 3E**, **top**), and a higher and sharper signal at the start codon in the HT samples (**Figure 3E**, **bottom**). Finally, the accuracy of individual uTLs detection was assessed by intersecting our datasets with a list of uTLs-derived peptides identified in proteomic datasets (Mudge et al., 2022). Even though substantial overlap is not expected, given the predominant function of uTLs in translation regulation rather than in encoding peptides (Ruiz-Orera et al., 2019; Orr et al., 2019), this analysis aimed to further assess the coding potential of identified uTLs. Remarkably, we observed that RiboWich captures a higher number of uTLs associated with detected peptides compared to RiboSeq (**Figure 3F** and **Supplementary Table 4**), confirming that our innovative technology outperforms existing RiboSeq protocols in capturing uTLs, with high-quality and at single-nucleotide resolution.

**Figure 3.**
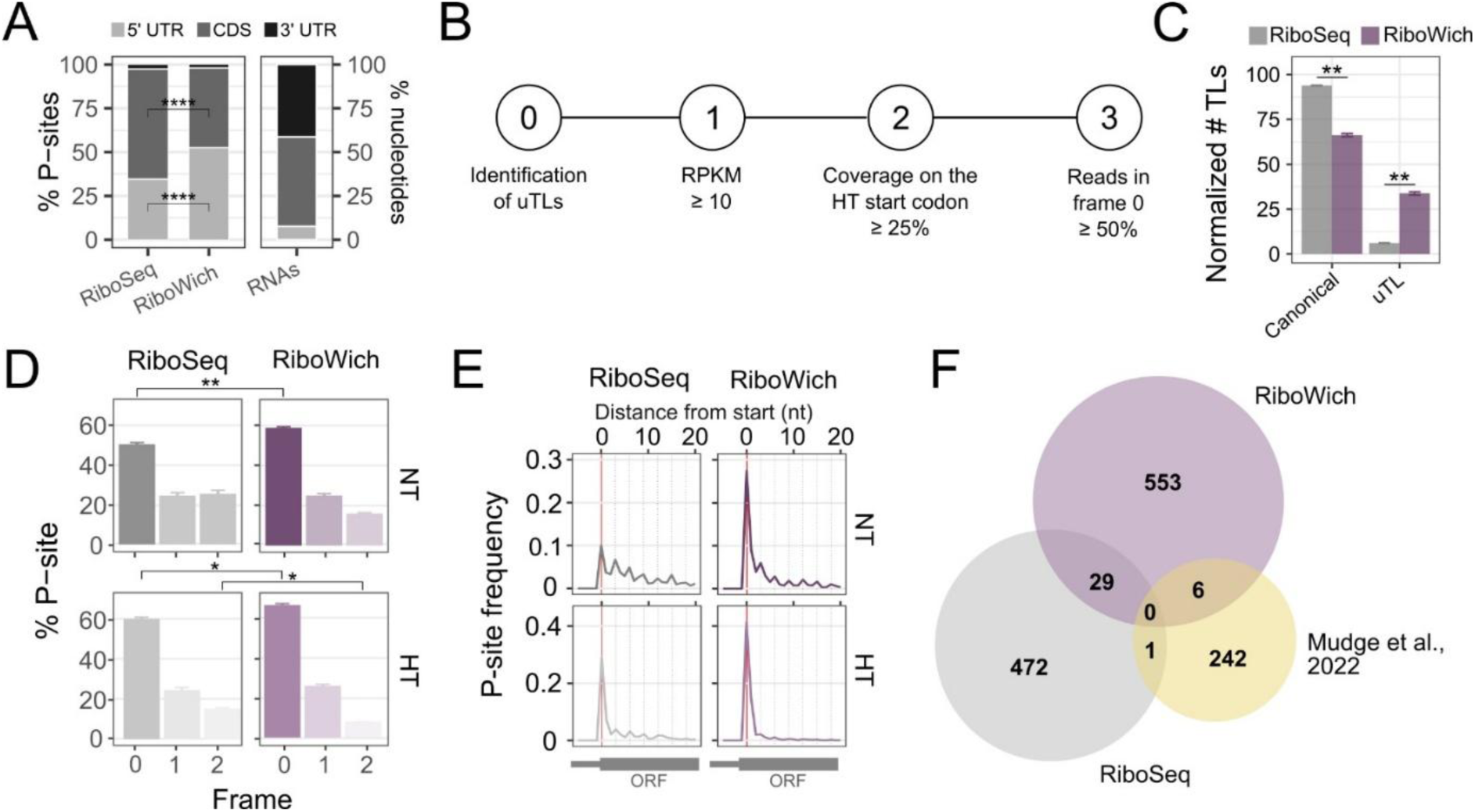
Upstream TLs in RiboWich and RiboSeq. (**A**) Percentages of P-sites mapping on the 5’ UTR, CDS, and 3’ UTR for RiboSeq HT (left) and RiboWich HT (middle). Percentages of region lengths in mRNA sequences are represented as reference (right). Results represent the mean of n=3 independent technical replicates. Two-sided T-test: P-value < 0.0001 (***). (**B**) Schematic representation of the workflow used to define active uTLs. (**C**) Percentage of active uTLs normalized against the total TLs (canonical and upstream). Results represent the mean ± SEM of n=3 independent technical replicates. Two-sided T-test: P-value < 0.01 (**). (**D**) Percentage of P-sites mapping to the three reading frames for RiboSeq (left) and RiboWich (right) NT (top) and HT (bottom) samples. Results represent the mean ± SEM of n=3 independent technical replicates. Two-sided T-test: P-value < 0.05 (*) and < 0.01 (**). (**E**) Meta-profiles showing the frequency of P-sites mapping at the beginning of the coding sequence for RiboSeq (left) and RiboWich (right) NT (top) and HT (bottom) samples. Results represent the mean ± SEM of n=3 independent technical replicates. (**F**) Intersection of the active uTLs identified in RiboSeq and RiboWich and the uTLs associated with at least one peptide detected in proteomics studies according to *Mudge et al., 2022*.

### RiboWich allows robust profiling in low-input samples

To expand on these findings and explore *in vitro* RiboWich (**Figure 1A**) in low-input samples, we prepared libraries using 25 femtomoles of purified ribosomes from non-treated (NT) and harringtonine-treated (HT) primary cortical neurons (CNs) and astrocytes from wild-type C57BL/6 mice. Usually, these samples pose significant challenges due to the low extraction yield of macromolecules, specifically ribosomes and RPFs (Verma et al., 2020). To optimize the processing of low-input samples, we enhanced the RiboWich workflow in two key areas. First, as proof of principle, we introduced multiplexed adaptors for barcoding REFs, enabling the simultaneous processing of six distinct samples. Following the ligation, the samples were pooled into a single tube, reducing sample loss and improving the efficiency of downstream enzymatic steps (**Figure 4A**). Second, we developed a gel-free REF preparation protocol, eliminating the need for gel-based size-selection and purification, significantly reducing the risk of material loss.

**Figure 4.**
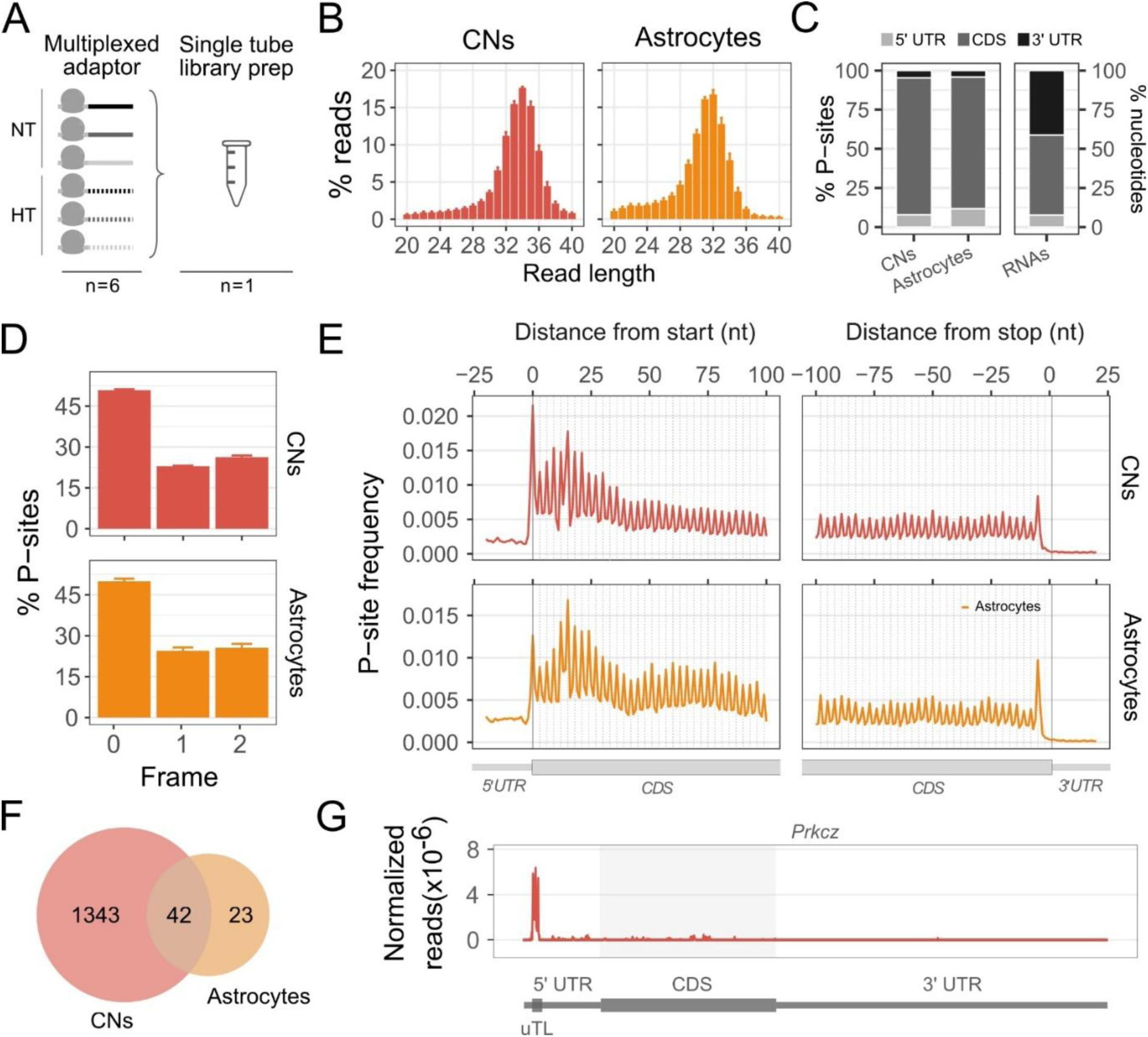
RiboWich in NT primary CNs and astrocytes. (**A**) Schematic representation of the RiboWich in multiplexing workflow (**B**) Percentage distribution of read lengths for CNs (left) and astrocytes (right). Results represent the mean ± SEM of n=2 (CNs) or n=3 (astrocytes) independent technical replicates. (**C**) Percentages of P-sites mapping on the 5’ UTR, CDS, and 3’ UTR for CNs (left) and astrocytes (middle). Percentages of region lengths in mRNA sequences are reported as reference (right). Results represent the mean of n=2 (CNs) or n=3 (astrocytes) independent technical replicates. (**D**) Percentage of P-sites mapping to the three reading frames for CNs (top) and astrocytes (bottom). Results represent the mean ± SEM of n=2 (CNs) or n=3 (astrocytes) independent technical replicates. (**E**) Meta-profiles showing the frequency of P-sites mapping at the beginning and the end of the coding sequence in CNs (top) and astrocytes (bottom). Results represent the mean ± SEM of n=2 (CNs) or n=3 (astrocytes) independent technical replicates. (**F**) Intersection of active uTLs detected in primary CNs and astrocytes. (**G**) Read coverage profiles for the CNs-specific *Prkcz* transcript, normalized for the library size for each replicate. Results represent the mean ± SEM of n=2 independent technical replicates.

The distribution of read lengths peaks within the expected range of RPF lengths (30-36 nt) for both cell types and in both conditions (**Figure 4B** and **Supplementary Figure 3A**). For both cell types in NT conditions, the majority of P-sites map to the CDS, with lower percentages mapping to the 5’ and 3’ UTRs (**Figure 4C**). As expected, the HT samples display a higher percentage of P-sites mapping at the 5’ UTR (**Supplementary 3B**). Next, we verified the trinucleotide periodicity of ribosome P-sites in the CDS of NT and HT samples (**Figure 4D-E** and **Supplementary 3C-D**). In line with the harringtonine function, the HT samples show a relative increase in the signal at the start codon and a decrease along the CDS (**Supplementary 3D**). Prompted by the observation that RiboWich captures *bona fide* RPFs in both cell types, we investigated the presence of uTLs in CNs and astrocytes following the previously described approach (**Figure 3B**). We identified 1,343 CNs-specific and 23 astrocyte-specific uTLs, and 42 uTLs shared between the two cell types (**Figure 4F** and **Supplementary Table 5**). However, differences in the absolute numbers of uTLs may be influenced by variations in sequencing depth between CNs and astrocytes. Interestingly, the uTL of *Prkcz*, included among the CNs-specific uTLs, is known to be associated with peptides detected in proteomics data (Mudge et al., 2022) and to regulate the translation of its canonical TL (Bal et al., 2016) (**Figure 4G**).

These results support our previous findings in HEK293T cells and highlight the key role of multiplexing in enabling efficient low-input sample processing without compromising data quality.

### RiboWich enables direct ligation to the REFs in low-input cellular lysates

After establishing the *in vitro* feasibility of RiboWich, we applied RiboWich in multiplexing directly to cellular lysates to evaluate its efficiency and versatility in a more relevant context. To further reduce input requirements, we adapted the workflow to a plate format.

To demonstrate the versatility of in-lysate RiboWich for low-input and plate-based applications, we carried out cell lysis coupled with RNase digestion, enzymatic inhibition, and adaptor ligation directly in the 96-well cell culture plate, where HEK293T cells (NT and HT) were previously seeded (**Figure 5A**). Critically, the library preparation was performed using two different inputs of either 20,000 (20K) or 2,000 (2K) cells per well. In both cases, the distribution of read lengths from NT samples peaks within the expected range of RPFs, indicating high library quality even when starting from very low input material (**Figure 5B**). For both 20K and 2K, most P-sites map to the CDS, with lower proportions mapping to the 5’ and 3’ UTRs (**Figure 5C**). Moreover, clear trinucleotide periodicity of ribosome P-sites is observed within the CDS in the two samples (**Figure 5D-E**). Importantly, this high-quality signal is maintained in HT-treated samples, both with 20K and 2K input cells (**Supplementary Figure 4A-D**).

**Figure 5.**
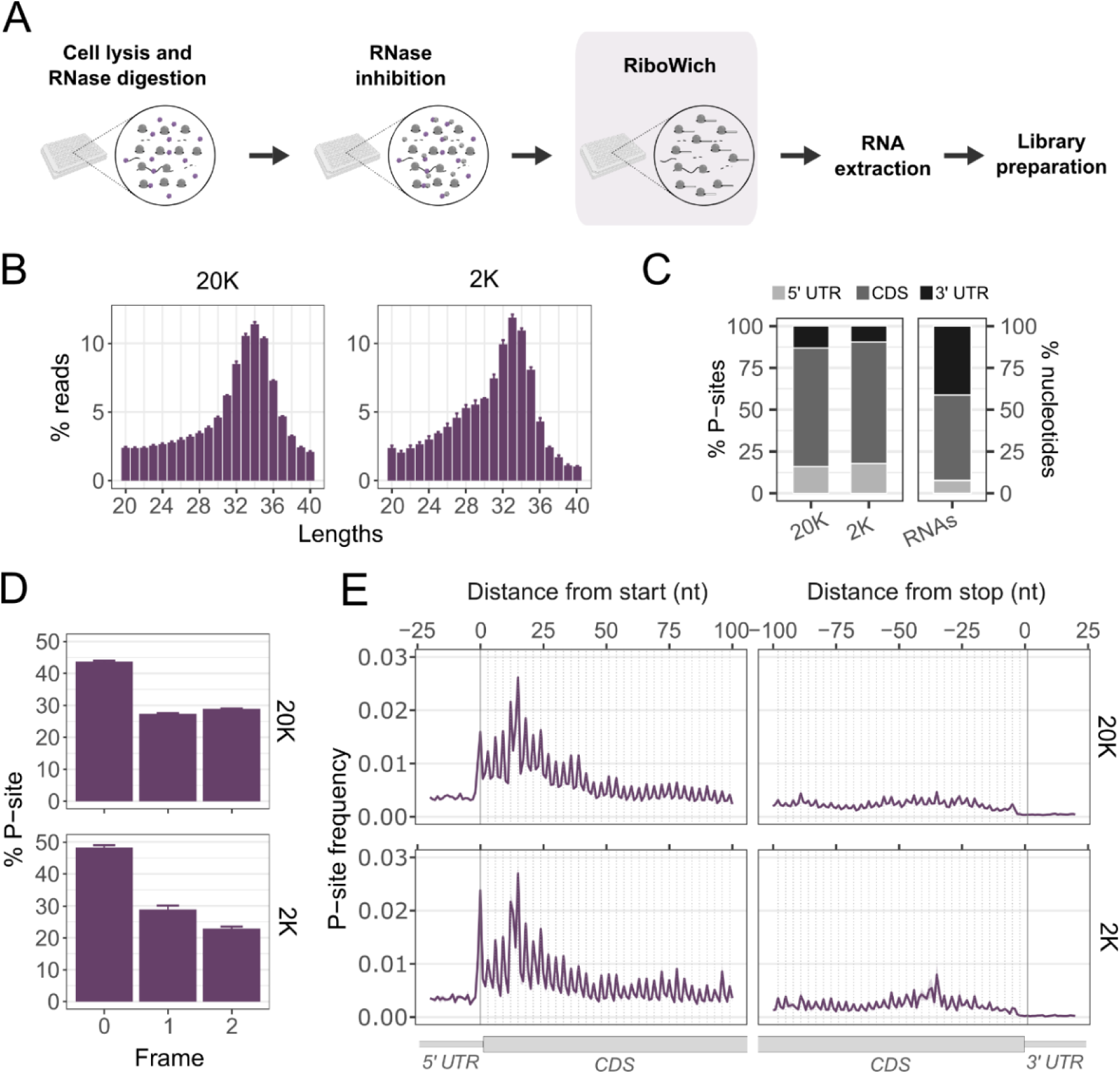
Low-input RiboWich: comparison between 20,000 (20K) and 2,000 (2K) HEK293T. (**A**) Experimental workflow. Cells are lysed and treated with RNase I. Upon nuclease inhibition, multiplexed adaptor ligation is performed directly in-lysate. RNA is extracted, and the following steps of library preparation are carried out in a single tube. (**B**) Percentage distribution of read lengths for 20K (left) and 2K (right). Results represent the mean ± SEM of n=6 (20K) or n=18 (2K) independent technical replicates. (**C**) Percentages of P-sites mapping on the 5’ UTR, CDS, and 3’ UTR for 20K (left) and 2K (middle). Percentages of region lengths in mRNA sequences are reported as reference (right). Results represent the mean of n=6 (20K) or n=18 (2K) independent technical replicates. (**D**) Percentage of P-sites mapping to the three reading frames for 20K (top) and 2K (bottom). Results represent the mean ± SEM of n=6 (20K) or n=18 (2K) independent technical replicates. (**E**) Meta-profiles showing the frequency of P-sites mapping at the beginning and at the end of the coding sequence in 20K (top) and 2K (bottom). Results represent the mean ± SEM of n=6 (20K) or n=18 (2K) independent technical replicates.

Together, these results show that in-lysate RiboWich is compatible with in plate processing and remains effective even with minimal input material.

### High-throughput RiboWich allows the simultaneous processing of 96 samples

Finally, to advance ribosome profiling towards high-throughput applications, we implemented a dual-step multiplexing RiboWich strategy. (**Figure 1B**). We seeded an entire 96-well plate with four diverse cell lines (one murine and three human) and subjected the cells to eight treatments, resulting in a total of 32 conditions, with three replicates each (**Figure 6A**).

**Figure 6.**
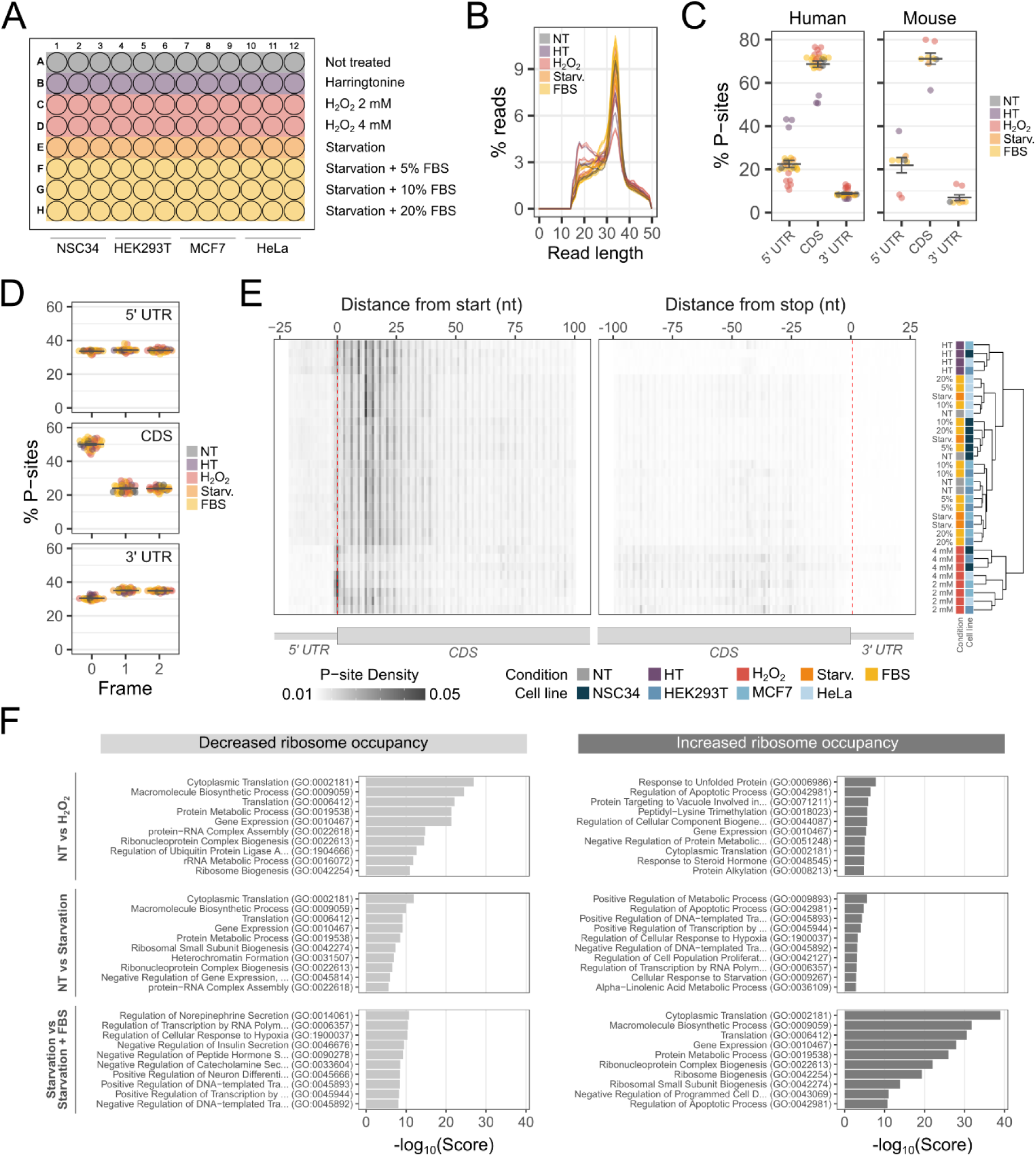
High-throughput RiboWich with 96 samples. (**A**) Schematic of the 96-well plate layout. (**B**) Percentage distribution of read lengths. Each line represents the mean of n=3 of technical replicates of each condition for each cell line. (**C**) Percentages of P-sites mapping on the 5’ UTR, CDS, and 3’ UTR, shown separately for human and mouse samples. Results represent the mean ± SEM of n=8 conditions for n=4 cell lines. Each dot represents the mean of n=3 technical replicates. Percentages of region lengths in mRNA sequences are reported in Figure 2C (human) and Figure 4C (mouse) as reference. (**D**) Percentage of P-sites mapping to the three reading frames. Results represent the mean ± SEM of n=8 conditions for n=4 cell lines. Each dot represents the mean of n=3 technical replicates. (**E**) Heatmap of meta-profiles showing the frequency of P-sites mapping at the beginning and at the end of the coding sequence. Each row represents the mean of n=3 technical replicates. Unsupervised clustering of genes was performed according to the signal reported. (**F**) Gene Ontology (GO) enrichment analysis of ribosome occupancy changes. The top 10 Biological Process terms are shown for Not-treated vs H₂O₂ stress, Not-treated vs Starvation, and Starvation vs Starvation + FBS. In panels (B-E), NT: not treated, HT: harringtonine-treated, H_2_O_2_: H_2_O_2_ 2 mM and 4 mM, Starv.: starvation, and FBS: starvation + 5%, 10%, 20% FBS.

For all conditions, the percentages of reads mapping to the transcriptome are comparable between all samples, suggesting no barcode bias (**Supplementary Figure 5A**). For all conditions, read length distributions show the expected peak for RPFs (**Figure 6B**), with only a modest increase in shorter reads for the H_2_O_2_ conditions, possibly reflecting ribosomal conformational changes under oxidative stress (Simms et al., 2014). In both mouse and human cell lines, the majority of P-sites map to CDS (**Figure 6C**). As expected, HT samples display a relative increase in the percentage of P-sites within 5′ UTRs. In contrast, H_2_O_2_-treated samples show a relative increase in the percentage of P-sites within 3′ UTRs, consistent with the notion that oxidative stress promotes translational readthrough beyond stop codons (Gerashchenko et al., 2012).

The overall data quality is further confirmed by the clear trinucleotide periodicity of P-sites, with more than 50% P-sites mapping to the correct frame in the CDS (**Figure 6D**). Notably, RiboWich resolves positional differences in ribosome occupancy between conditions, as illustrated by the meta-profiles (**Supplementary Figure 5B**) and the collapsed heatmap of meta-profiles (**Figure 6E**). HT samples display the expected accumulation of ribosome signal at the start of the CDS, which progressively decreases toward the end. In contrast, H_2_O_2_-treated samples show reduced ribosome density at the beginning of the CDS and increased signal toward the end, in line with previous works reporting altered elongation dynamics under oxidative stress (Shan et al., 2007; Sanchez et al., 2019). To further explore the data, we examined ribosome occupancy under oxidative stress and starvation compared to control samples (**Supplementary Figure 5C** and **Supplementary Table 6-7**). The most pronounced changes are related to the H_2_O_2_-treated samples, where, across all 4 cell lines, the vast majority of transcripts displayed decreased ribosome occupancy compared to NT, rather than an increase. Finally, to gain further insights, we performed Gene Ontology (GO) enrichment analysis on these differentially occupied transcripts (**Figure 6F** and **Supplementary Table 8-9**). Both oxidative stress- and starvation-treated samples exhibit reduced ribosome occupancy on translation-related genes, and this effect is reversed when starved cells are re-supplemented with FBS. Conversely, stress-specific responses, including unfolded protein response and apoptotic processes, are enriched under both stress conditions, and these terms are lost when starvation is rescued.

Overall, RiboWich enables high-throughput, low-input ribosome profiling across multiple cell lines and conditions, accurately capturing both global and transcript-specific translational changes while resolving positional differences in ribosome occupancy.

## Discussion

Over the past 15 years, ribosome profiling has revolutionized our ability to investigate protein synthesis across a wide range of biological systems (Ingolia et al. 2009). Yet, despite its widespread use, traditional ribosome profiling methods face several limitations that may impact data quality (Wang and Mao, 2023). Such limitations include i) time-consuming protocols, ii) difficulties in obtaining high-quality data from challenging, low-input samples, and iii) the inability to process a high number of samples simultaneously (Wang and Mao, 2023; Tomuro and Iwasaki, 2025). Together, these obstacles hinder the development of ribosome profiling in low-input, high-throughput, single-cell, and spatial translatomics applications, highlighting the urgent need for audacious technological innovations.

To overcome these challenges, we developed RiboWich (<u>Ribo</u>some sand<u>Wich</u>), a first-of-its-kind technology that enables the direct ligation of sequencing adaptors to RNA fragments whilst embedded within ribosomes. This innovative and unique approach bypasses the need for ribosome isolation, RNA extraction, RPF size selection, and RPF purification, thereby simplifying and accelerating the workflow. As proof of principle, we demonstrated that *in vitro* RiboWich efficiently captures authentic RPFs, the length of which is in line with the length of *bona fide* ribosome footprints, enriched within coding sequences and maintaining the correct reading frame, providing robust ribosome positional information in both cell lines and primary cells. These findings underscore the strength and reliability of RiboWich as a valuable method for ribosome profiling, offering elevated efficiency and data quality for studying protein synthesis.

The non-optimal signal-to-noise ratio often observed in RiboSeq experiments hampers the detection and analysis of non-canonical translational events (Mao and Qian, 2023), which, despite occurring at low frequency, play crucial roles in nuanced translational regulation, particularly in stress responses (Khitun et al., 2019; Moro et al., 2021), cellular differentiation (Vatikioti et al., 2019), and various disease contexts (Silva et al., 2019). We demonstrated that RiboWich detects a significantly greater number of active uTLs, thus surpassing the capabilities of the standard ribosome profiling method. We also observed that RiboWich captures a higher number of uTLs associated with detected peptides (Mudge et al., 2022) compared to RiboSeq. Moreover, we identified CNs- and astrocytes-specific uTLs that allow an accurate dissection of translation differences between the two cell types. These results provide strong evidence that RiboWich unravels active translation occurring in non-canonical TLs, offering a more comprehensive and accurate approach to studying translation regulation.

While recent studies have made great strides toward low-input ribosome profiling, generating libraries from as few as thousands of cells (Hornstein, N. et al., 2016; Xiong et al., 2022; Li et al., 2022; Zhang et al.,2022; Meindl et al., 2023; Froberg et al.,2023; Ozadam et al., 2023; summarized in **Table 1**) or from single cells (VanInsberghe et al., 2021), these methods come with specific limitations tied to the library construction itself. These techniques rely on the template-switch or the dual-ligation method, which, while powerful, have sequence biases and are not compatible with early-stage barcode integration, making them unsuitable for multiplexing (Wang and Mao, 2023). From a technical perspective, RiboWich introduces several innovations that collectively streamline ribosome profiling and extend its applicability to previously challenging contexts. By performing adaptor ligation directly in cellular lysates, the method eliminates the need for ribosome purification, shortens the workflow, and preserves material from low-input samples, while remaining fully compatible with ribosome isolation strategies such as polysome profiling or active ribosome isolation (Clamer et al., 2018), which can be applied after ligation. The incorporation of early barcoding enables efficient multiplexing, allowing multiple samples to be pooled immediately after ligation and processed in a single tube throughout all downstream steps. This strategy minimizes material loss and reduces the time and costs typically associated with processing each sample independently. Finally, when implemented in plate-based formats, RiboWich enhances scalability and throughput, making it an ideal solution for interrogating dozens of conditions, replicates, or time points in a single experiment. Thus, RiboWich is ideal for processing extremely low-input samples, as we successfully demonstrated profiling as few as 2,000 cells. Together, these innovations establish RiboWich as a versatile, high-performance platform that bridges the gap in low-input ribosome profiling.

**Table 1.**
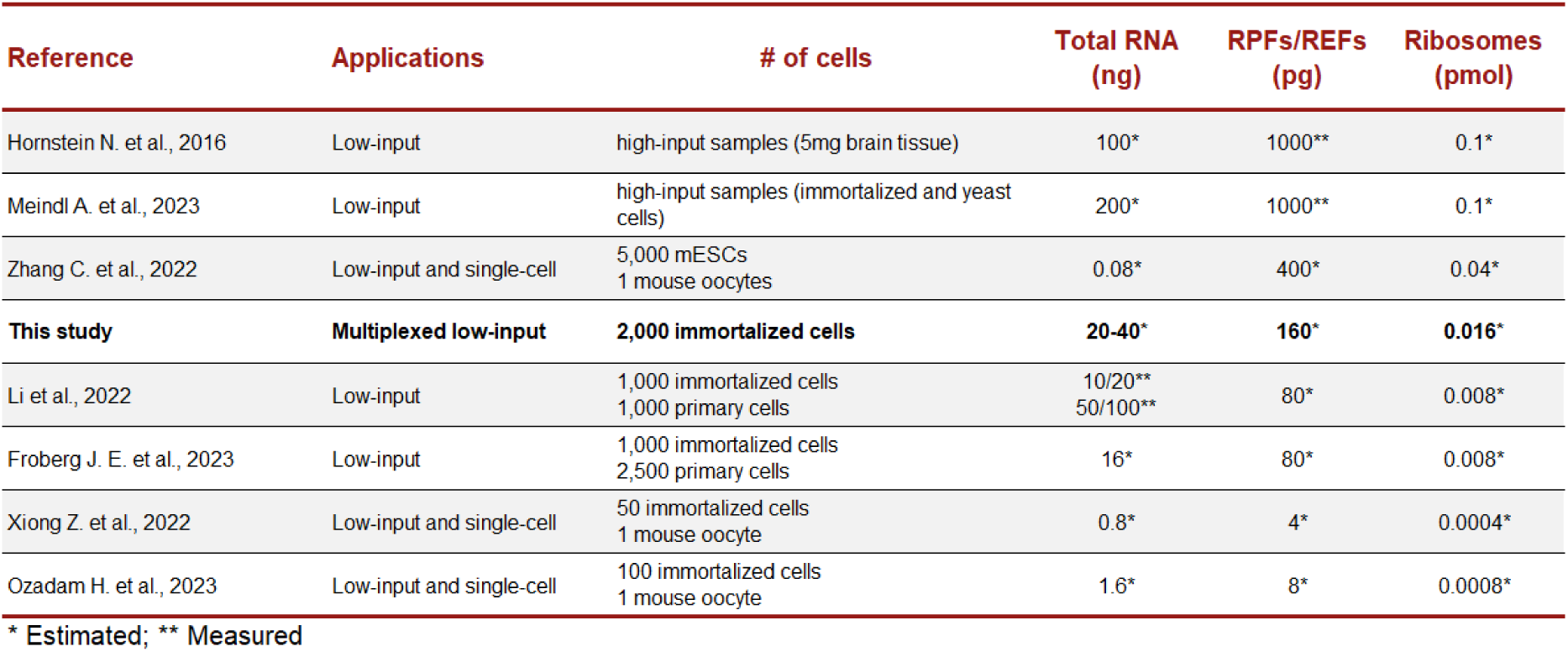
Comparison of low-input ribosome profiling studies.

| Reference | Applications | # of cells | Total RNA (ng) | RPFs/REFs (pg) | Ribosomes (pmol) |
| --- | --- | --- | --- | --- | --- |
| Hornstein N. et al., 2016 | Low-input | high-input samples (5mg brain tissue) | 100* | 1000** | 0.1* |
| Meindl A. et al., 2023 | Low-input | high-input samples (immortalized and yeast cells) | 200* | 1000** | 0.1* |
| Zhang C. et al., 2022 | Low-input and single-cell | 5,000 mESCs<br>1 mouse oocytes | 0.08* | 400* | 0.04* |
| **This study** | **Multiplexed low-input** | **2,000 immortalized cells** | **20-40\*** | **160\*** | **0.016\*** |
| Li et al., 2022 | Low-input | 1,000 immortalized cells<br>1,000 primary cells | 10/20**<br>50/100** | 80* | 0.008* |
| Froberg J. E. et al., 2023 | Low-input | 1,000 immortalized cells<br>2,500 primary cells | 16* | 80* | 0.008* |
| Xiong Z. et al., 2022 | Low-input and single-cell | 50 immortalized cells<br>1 mouse oocyte | 0.8* | 4* | 0.0004* |
| Ozadam H. et al., 2023 | Low-input and single-cell | 100 immortalized cells<br>1 mouse oocyte | 1.6* | 8* | 0.0008* |

\* Estimated; \*\* Measured

Despite some progress in scalable translatomic analysis, no method has yet enabled high-throughput ribosome profiling. Recently, riboPLATE-seq introduced a scalable approach to link ribosome association with transcript quantification, but its resolution remains limited as it does not capture codon-level dynamics (Metz et al., 2022). The high-throughput RiboWich approach utilizes a dual-step multiplexing strategy, introducing unique RNA barcodes during the first ligation and distinct DNA barcodes during reverse transcription. This design enables scalable multiplexing, allowing the simultaneous processing of a large number of samples. Consequently, the workflow minimizes hands-on time, requiring only about two days from cell lysis to sequencing-ready libraries, and drastically reduces the cost per sample. RiboWich also promotes standardization and reproducibility, making it ideal for high-throughput screening. We demonstrated its robustness by processing 96 samples in parallel, obtaining high-quality and consistent data across all replicates. Crucially, RiboWich captures condition-specific translation signatures, as evidenced by reproducible responses to harringtonine, oxidative stress, and nutrient deprivation, underscoring its biological relevance. Together, these features establish RiboWich as the first and currently only method enabling true high-throughput ribosome profiling, uniquely combining technical simplicity with the sensitivity and resolution required for robust translational studies.

Several aspects of RiboWich deserve further consideration. The method tends to capture a broader distribution of read lengths compared to conventional RiboSeq. This outcome is expected, as RiboWich omits the size-selection step and therefore retains a wider spectrum of ribosome-protected RNA fragments. Moreover, the number of ribosomes per cell can vary substantially between cell types, directly influencing the amount of material available for library preparation. Consequently, RiboWich may still require sample-specific optimization when handling thousands of cells from challenging samples to ensure efficient library preparation and the generation of robust translatome data.

In conclusion, RiboWich sets a new paradigm for ribosome profiling by simplifying workflows, improving data quality, and enabling true low-input, high-throughput applications such as drug screening. By overcoming long-standing technical bottlenecks with robust multiplexing, it positions ribosome profiling alongside other high-throughput omics, ensuring translation research keeps pace. Most importantly, RiboWich provides the scalable and reproducible data foundation needed to integrate the translatome into large-scale platforms and fuel AI-driven discoveries, while paving the way toward single-cell and spatially resolved translation investigations, the next frontier in precision medicine.

## Materials and Methods

### Cell lines

HEK293T, MCF7, HeLa, and NSC34 cell lines were maintained in culture in Dulbecco’s Modified Eagle’s Medium (DMEM, Euroclone), 10% FBS (GIBCO, Life Technologies), 2 mM L-Glutamine (GIBCO, Life Technologies), and 1% penicillin/streptomycin (GIBCO, Life Technologies) at 37°C in a 5% CO_2_ atmosphere.

### Wild-type C57BL/6 mice for primary cortical neurons and astrocytes cultures

Wild-type C57BL/6 mice were obtained from Charles River and were used to prepare primary cortical neurons and astrocytes cultures. Animal care and experimental procedures were performed in accordance with the Ethical Committee of the University of Trento and were approved by the Italian Ministry of Health (576/2021-PR). Animals were maintained with access to food and water ad libitum and kept at a constant temperature (19–22°C) with a 12:12 h light/dark cycle.

### Cultures of primary cortical neurons from wild-type C57BL/6 mice

Primary cortical neurons were obtained from wild-type mice at embryonic stage E15.5 as previously described (Migazzi et al., 2021). Meninges were removed from cortices and were digested in papain solution (20U papain, 500 µM EDTA, 100 µM Cystine, 26 mM Sodium Bicarbonate in 1X Earle’s Balanced Salt Solution (EBSS, Gibco) at 37°C for 20 minutes. After DNase I treatment at 37°C for 3 minutes, they were centrifuged at 1578 rcf for 5 minutes. The pellet was resuspended in EBSS supplemented with Bovine Serum Albumin and Trypsin Inhibitor. Cells were centrifuged at 1578 rcf for 10 minutes and resuspended in the plating medium (10% FBS, 1% Pen Strep in MEM with L-Glutamine). Cortical neurons were seeded on pre-coated wells with poly-D-lysine (Sigma). The day after, the whole medium was replaced with Neurobasal medium added with 1% B27, 1 mM Sodium Pyruvate, 1% penicillin/streptomycin, 2 mM L-Glutamine, and 8 µM AraC (Sigma). Cortical neurons were cultured at 37°C with 5% CO_2_.

### Cultures of astrocytes from wild-type C57BL/6 mice

Primary astrocytic cells were cultured from wild-type newborn pups (P0-P2). After meninges removal, cortical tissues were enzymatically digested in papain solution and treated with DNase I, following the same procedure as for neuronal culture. After digestion blockage, cells were resuspended in DMEM (Gibco) supplemented with 10% FBS, 2 mM L-Glutamine, and 1% penicillin/streptomycin and plated. At DIV1, the medium was replaced with fresh complete medium. Cells were cultured at 37°C with 5% CO_2_.

### RiboSeq

Cytoplasmic lysates were prepared according to (Tebaldi et al., 2012). Briefly, cells were seeded in 10 cm diameter dishes and grown until they reached 70-80% confluence. Specifically for the harringtonine-treated (HT) samples, cells were treated for 10 minutes with 2 µg/mL harringtonine (Abcam, ab141941-10) at 37°C. Then, not-treated (NT) and HT cells were incubated for 3 minutes and 30 seconds at 37°C with cycloheximide (10 μg/mL) (Sigma Aldrich, C7698), an inhibitor of translation elongation that was added to the culture medium to stabilize ribosomes on mRNAs. Upon incubation, dishes were put on ice and washed three times with cold PBS (Gibco, Life Technologies) supplemented with 10 μg/mL cycloheximide. Next, 100-300 μL of the Standard Cytoplasmatic Lysis Buffer (10 mM Tris–HCl pH 7.5, 10 mM MgCl2, 10 mM NaCl, 1% Triton X-100, 5 U/mL DNase I (Thermo Scientific), 0.2 U/µL RiboLock RNase Inhibitor (Thermo Scientific), 1 mM DTT (Sigma Aldrich), 10 µg/mL cycloheximide, 1% Na-deoxycholate) were added depending on the experiment, and cells were lysed with the help of a scraper. The lysates were collected and cleared from cellular debris by centrifugation for 5 minutes at 14,000 g at 4° C. The supernatants were transferred into new tubes, and the RNA content of each lysate was assessed by measuring the absorbance at 260 nm using Thermo Scientific NanoDrop. Next, since lysates were obtained in hypotonic conditions, the NaCl concentration was adjusted to 100 mM. If not processed immediately, samples were stored at -80 °C.

For RNase I digestion, polysome profiling, and RNA extraction, lysates were treated with RNase I (2/3 Invitrogen AM2295 and 1/3 BioResearch Technologies N6901K) using 7.5 U / a.u. Abs260 of lysate, and incubated for 45 minutes on an orbital shaker at RT. The RNase I activity was inhibited by the addition of 100U of SUPERase·In™ RNase Inhibitor (Invitrogen, AM2696). Samples were kept on ice for 10 minutes. In parallel, the sucrose gradient was prepared in cold 13.2 mL polyallomer ultracentrifuge tubes (Beckman) pouring 5.5 mL of 40% w/V sucrose (in 100 mM NaCl, 10 mM MgCl_2_, 10 mM Tris/HCl pH 7.5) at the bottom of the tube which was then gently filled with 10% w/V sucrose (in 100 mM NaCl, 10 mM MgCl2, 10 mM Tris/HCl pH 7.5). The tubes were slowly tilted to a horizontal position and maintained for 2 hours at 4°C to allow the gradient formation. After 2 hours, they were placed in a vertical position and kept on ice. From the top, 700 μL of sucrose were removed, and the cytoplasmic lysate was gently loaded drop by drop. The samples were ultracentrifuged for 1 hour and 30 minutes at 40,000 rpm at 4 °C in the Beckman Optima XPN-100 Ultracentrifuge using the SW41 rotor. After ultracentrifugation, gradients were fractionated using a Teledyne ISCO model 160 fractionator equipped with a UA-6 UV/ VIS detector to measure absorbance at 254 nm. For each sample, 12 fractions were collected, each of 1000 μL in volume. Fractions corresponding to the 80S monosome were collected, treated with 100 μL/mL of 10% SDS and 5 μL/mL of Proteinase K (PanReac AppliChem, #A4392), and incubated at 37 °C for 1 hour and 30 minutes. Next, 250 μL/mL of Phenol:Chloroform:Isoamyl Alcohol (Sigma, P3803) were added, and the samples were centrifuged for 10 min at 10,621 g at 4°C. The upper aqueous phase was transferred into a new tube, and RNA was precipitated in 1 mL of isopropanol with the addition of 1 μL of GlycoBlue Coprecipitant (Invitrogen, AM9516) for each sample. Upon precipitation at -80 °C overnight, samples were centrifuged at 15,294 g for 45 minutes at 4°C. The pellets were washed with 500 μL of cold 80% ethanol, and, after complete drying, RNA was solubilized in 5 μL of nuclease-free water.

For size selection and PAGE purification of RPFs, RNA from each sample was mixed with the 2x denaturing TBE-Urea Sample Buffer (Invitrogen, LC6876) and denatured at 80 °C for 90 seconds. RNA was run on a 15% TBE-Urea gel (Invitrogen, EC6885BOX) for 1 hour and 10 minutes at 170V. After fast staining with SYBR Gold Nucleic Acid Gel Stain (Invitrogen, S11494), the gel was visualized using Bio-Rad Chemidoc. The fragments between 28-32 nt were excised, and the band was dissolved in 400 μL of nuclease-free water, 40 μL of ammonium acetate 5M, and 2 μL of 10% SDS using a pestle. Samples were left overnight at 4 °C in rotation. The next day, they were transferred onto Ultrafree Centrifugal Filter Units 0.22 μm (Millipore, UFC30GV0S) and centrifuged according to the manufacturer’s protocol. The RNA was precipitated in 700 μL of isopropanol, as previously described, and finally resuspended in 20 μL of nuclease-free water.

For the library preparation with the standard RiboSeq, 40 ng of RPFs underwent the steps of sequencing library preparation according to the Immagina Biotechnology protocol for LaceSeq^TM^ (LS001-12). Upon amplification, libraries were purified, as previously described, using 8% TAE gel electrophoresis (40 minutes at 150V), and the dsDNA was solubilized in 12 μL of nuclease-free water. Their quality was assessed using High Sensitivity DNA CHIPS (Agilent Technologies, #5067-4626) on the Agilent 2100 Bioanalyzer. Eventually, libraries were analyzed by single-read sequencing 100 cycles with Illumina NovaSeq 6000 by the Next Generation Sequencing Facility (CIBIO, University of Trento, Italy).

### In vitro RiboWich

Cytoplasmatic lysates from NT and HT HE293T cells, cortical neurons (CNs), and astrocytes were prepared as previously described. RNase I treatment, nuclease inhibition, and polysome profiling were performed as previously illustrated. Ribosomes were purified from fractions corresponding to the 80S monosome by ultracentrifuging for 1 hour and 30 minutes at 100,000 rpm at 4 °C in the Benchtop Beckmann Ultracentrifuge, Optima TLX ultracentrifuge using the TLA100.2 rotor. Supernatants were discarded, and ribosomes were resuspended in 10-40 μL of the *in vitro* Ribosome Solution (75 mM KCl, 50 mM Tris-HCl pH 8.1, 5 mM MgCl2 in DEPC, freshly supplemented with 10 mM DTT (Sigma Aldrich), 10 µg/mL cycloheximide, 0.2 U/µL RiboLock RNase Inhibitor (Thermo Scientific)). Ribosome concentrations were determined by measuring the absorbance at 260 nm with the Thermo Scientific NanoDrop and employing the published ribosome extinction coefficient (5×10^7^ cm^-1^ M^-1^) (Algire et al., 2002). Ribosomes were eventually stored at -80°C, after fast freezing in liquid nitrogen.

For the *in vitro* RiboWich experiment in HEK293T, ribosomes were incubated for 2 hours at 37 °C in rotation with the *in vitro* RiboWich Ligation Solution (*in vitro* Ribosome Solution supplemented with the sequencing adaptor (LaceSeq^TM^, LS001-12) in a ratio 1:2 (adaptor: ribosomes), 0.15 mM GTP (Invitrogen, AM8130G), 1.8 mM MnCl_2_, and 1 μL of Rtcb-like enzyme (Immagina Biotechnology srl). For the *in vitro* multiplexed RiboWich experiment in CNs and astrocytes, ribosomes were incubated for 2 hours at 37°C in rotation with the *in vitro* RiboWich Solution with multiplexed sequencing adaptors (6 distinct barcoded adaptors, Immagina Biotechnology Multiplexed LaceSeq^TM^, MX001-36) in a ratio 1:4 (adaptor: ribosomes), and the rest of the reagents as described above.

Upon incubation, RNA was extracted either with the TRIzol reagent (Thermo Fisher Scientific, 15596026) following the manufacturer’s protocol (gel approach) or with RNA Clean & Concentrator™-5 (Zymo Research, R1013, R1014, R1015, or R1016) following manufacturer’s instructions for small RNA purification (gel-free approach), respectively, for HE293T, and CNs and astrocytes. For the HEK293T experiment (gel approach), upon extraction, samples were resuspended in 5 μL of nuclease-free water and run on a 15% TBE-Urea gel (Invitrogen, EC6885BOX) and purified as previously described. The RNA was then resuspended in 20 μL of nuclease-free water. For the CNs and astrocytes experiment (gel-free approach), RNA was extracted, purified via columns, and finally eluted in 20 μL of nuclease-free water. In both abovementioned cases, the library preparation continued with the steps of 5’ phosphorylation, circularization, reverse transcription, and PCR amplification (according to the Immagina Biotechnology protocol for LaceSeq^TM^ (LS001-12)). Libraries were finally purified, quantified, and sent for sequencing as previously described.

### Low input RiboWich

HEK293T cells were seeded in 96-well plates and grown until they reached 40-60% or 5-10% (respectively for 20K and 2K) confluence. Cells for NT and HT conditions were treated as previously described. Upon cycloheximide incubation, the medium was removed, and cells were washed twice with the Preserving Solution (PBS supplemented with 100 µg/mL of cycloheximide). Next, 25 μL of RiboWich Cytoplasmatic Lysis Buffer (10 mM Tris–HCl pH 8.1, 5 mM MgCl_2_, 10 mM NaCl, 1% Triton X-100, 5 U/mL DNase I (Thermo Scientific), 0.4 U/µL RiboLock RNase Inhibitor (Thermo Scientific), 1 mM DTT (Sigma Aldrich), 100 µg/mL cycloheximide, 1% Na-deoxycholate) supplemented with 0.2 U/μL of RNase I (Invitrogen, AM2295) were added onto each well. The plate was incubated for 45 minutes on Mixer HC (Starlab) at 400 rpm at RT. The RNase I activity was inhibited by the addition of 1 U/µL of SUPERase·In™ RNase Inhibitor (Invitrogen, AM2696). The plate was incubated for 10 minutes on Mixer HC (Starlab) at 400 rpm at 37°C. Next, the lysates were adjusted to the in-lysate RiboWich Ligation Solution: Tris–HCl pH 8.1 and DTT concentrations were adjusted to 50 mM and 10 mM, respectively, 0.15 mM GTP (Invitrogen, AM8130G), 1.8 mM MnCl_2_, and 0.015 µM multiplexed sequencing adaptors (6 distinct barcoded adaptors, Immagina Biotechnology Multiplexed LaceSeq^TM^, MX001-36) were added together with 0.5 μL of Rtcb-like enzyme (Immagina Biotechnology srl), reaching a final volume of 30 μL per well. The plate was incubated for 2 hours on Mixer HC (Starlab) at 37°C at 400 rpm. Upon the ligation, differently barcoded samples from different wells were pooled, and RNA was directly extracted using the RNA Clean & Concentrator™-5 kit (Zymo Research, R1013, R1014, R1015, or R1016) following the protocol for Small RNAs and the manufacturer’s instructions. The library preparation continued with the phosphorylation at the 5’ end, circularization, reverse transcription, and PCR amplification according to Immagina Biotechnology’s protocol for LaceSeq^TM^ (LS001-12). Libraries were finally purified, quantified, and sent for sequencing as previously described.

### High throughput RiboWich

HEK293T, MCF-7, HeLa, and NSC-34 cells were seeded in 96-well plates and grown until reaching 40-60% confluence. Within the plate, all four cell types were subjected to the same experimental conditions: NT; treatment with HT as previously described, used at 10 µg/mL; treatment with H₂O₂ at 2 or 4 mM for 1 hour; starvation with 0% FBS for 16 hours, followed by re-addition of 5%, 10%, or 20% FBS for 1 hour (with one group maintained in starvation as control) (**Figure 6A**). After all the treatments, cells were treated with cycloheximide using a concentration of 100 μg/mL. In plate in-lysate RiboWich was performed as previously described up to the ligation step, using 8 distinct barcoded adaptors (Immagina Biotechnology Multiplexed LaceSeq^TM^, MX001-36). Samples from different wells, each carrying distinct barcodes, were pooled and purified using the RNA Clean & Concentrator™-5 kit (Zymo Research, R1013, R1014, R1015, or R1016), following the manufacturer’s instructions for small RNAs. Phosphorylation, circularization, and reverse transcription were carried out according to Immagina Biotechnology’s LaceSeq™ protocol (LS001-12). For reverse transcription, 12 distinct barcoded RT primers were used, with a distinct primer added to each reaction. Reactions were subsequently pooled into a single tube, and cDNA was precipitated using Na acetate 3M and isopropanol. PCR amplification was performed according to Immagina Biotechnology’s protocol for LaceSeq^TM^ (LS001-12). Libraries were finally purified, quantified, and sent for sequencing as previously described.

### NGS data analysis

For RiboSeq, RiboWich, and multiplexed RiboWich (CNs, astrocytes, and 20K/2K HEK293T), the single adaptors were removed with CutAdapt v3.7, and UMIs, composed of the first 4 and last 4 nucleotides of the remaining sequences, were extracted using UMI-Tools v1.1.6. For multiplexed RiboWich, the barcodes (spanning nucleotides 5 to 12) were extracted during the same step. For high-throughput RiboWich, the two barcodes, the UMIs and the sequences corresponding to the ribosome protected fragments were detected and extracted using UMI-Tools v1.1.6 using the following regex:

*^(?P<umi_1>.{4})(?P<cell_1>.{8}).+(?P<umi_2>.{4})(?P<discard_1>TCTCCTTGCAT AATCACCAACC){s<=4}(?P<cell_2>.{8})(?P<umi_3>.{4})(?P<discard_2>.+)*.

Next, the first nucleotide was trimmed, and reads shorter than 15 nucleotides were discarded. Reads mapping on rRNA and tRNA sequences were removed, and the remaining reads were mapped on the reference genome (human: GRCh38, ENSEMBL release 109 and Gencode v43; mouse: GRCm39, ENSEMBL release 109 and Gencode vM32 gene annotation) using STAR v2.7.10a. Downstream positional analyses of the reads mapping on the transcriptome were performed using riboWaltz (Lauria et al., 2018). For all the analyses, reads between 20-40 nt were selected, except for the high-throughput RiboWich, where reads between 32-35 nt were selected.

### Annotation of upstream TLs

Non-canonical uTLs were defined as sequences in the transcriptome with a start codon (either AUG or one of the 4 most common alternatives: CUG, UUG, GUG, ACG) detected before the canonical one, a stop codon (UAA, UAG, UGA) in frame with respect to the start site, a minimum of 6 nucleotides to include both the start and stop codons, and a length multiple of 3 nucleotides.

### Identification of upstream TLs

uTLs in RiboSeq and RiboWich were identified using a four-step filtering process based on features characteristic of canonical TLs. uTLs meeting the following criteria were retained: (i) RPKM > 10; (ii) at least 25% of total reads mapping to the start site in the HT condition; (iii) over 50% of total reads in the correct frame in the NT condition; and (iv) criteria (i)-(iii) satisfied in both RiboSeq and RiboWich datasets. The identified TLs were cross-referenced with the RiboUTL database (Liu, Qi, et al., 2023), obtained from https://rnainformatics.org.cn/RiboUORF. Additionally, the list of 274 uTLs associated with at least one peptide identified in proteomic datasets was sourced from Mudge et al., 2022 (Mudge et al., 2022).

### Differential analysis

Only protein-coding genes were retained for differential analyses. To address redundancy caused by reads mapping to distinct exons that span overlapping regions of the chromosomes, the exons of all isoforms of each gene were collapsed. Reads mapping to identical positions, except for one, were discarded. The number of P-sites from the remaining reads was then used to compute gene-specific coverage. To account for differences in library size and composition, gene coverages were normalized across replicates using the trimmed mean of M-values (TMM) method implemented in the edgeR R package. FPKMs were then calculated based on the length of the sequence resulting from the collapsed exons. Only genes with an FPKM ≥ 10 in all replicates of at least one condition were retained. Differential gene coverage analysis was performed using the glmQLFTest function from edgeR, applying a dual threshold to define genes showing significant changes in ribosome occupancy: (i) absolute log2 fold change ≥ 0.5 and (ii) adjusted p-value ≤ 0.05.

### Gene Ontology enrichment analysis

Gene Ontology (GO) enrichment analysis was performed using the EnrichR R package.

For high-throughput RiboWich, comparisons were grouped into three main categories: NT vs H₂O₂, NT vs Starvation, and Starvation vs Starvation and FBS. Enriched GO biological process terms were ranked by False Discovery Rate (FDR) within each category. To integrate the results, Robust Rank Aggregation (Kolde et al., 2012) was applied to the ranked lists, and the top 10 GO terms per category were selected for visualization.

## Data availability

RNA-Seq data generated by the current study have been deposited in the Gene Expression Omnibus (GEO) under the accession code [data in submission]. All other data supporting the findings of this study are available from the corresponding author upon reasonable request. Source data are provided in this paper.

## Acknowledgments

We thank the staff at the Core Facilities Next Generation Sequencing Facility (NGS) at the Department of CIBIO, University of Trento, for their technical support. We thank IMMAGINA Biotechnology, Italy, for funding this work and I.B., the European Union - Next Generation EU, Mission 4, Component 2, CUPB93D21010860004 A for supporting F.L.; the EU funding within the MUR PNRR “National Center for Gene Therapy and Drugs based on RNA Technology” (Project no. CN00000041 CN3 RNA) and the Caritro Foundation Bando ricerca e sviluppo 2021/2022 for supporting E.P. We are grateful to the Core Facilities Next Generation Sequencing Facility (NGS) at Department CIBIO University of Trento, Italy; and to Michela Laferla, Francesca Spanò and Elena Gerola (Administrative staff at IBF-CNR) for supporting with administrative management, and to Marta Marchioretto for her valuable help as Lab Manager at IBF-CNR.

## Author contributions

G.V. conceived and designed the study and directed the research. I.B., C.P., and E.P. designed and performed the experimental work. M.S., A.D.P., and M.C. designed the multiplexed adaptors. L.D. and M.B. prepared the murine primary cells. G.T. and F.L. processed all sequencing data and performed the analyses. G.V. obtained the funding. I.B., G.T., C.P., G.S., F.L., and E.P. prepared the figures. All authors contributed to the writing of the manuscript.

