## Supplementary File for "Empowering multiplexed ultra-throughout ribosome profiling with RiboWich"

[1] IMMAGINA Biotechnology s.r.l. (Italy)
[2] Institute of Biophysics, CNR Unit at Trento, (Italy)
[3] University of Trento (Italy)

* These authors equally contributed to this work

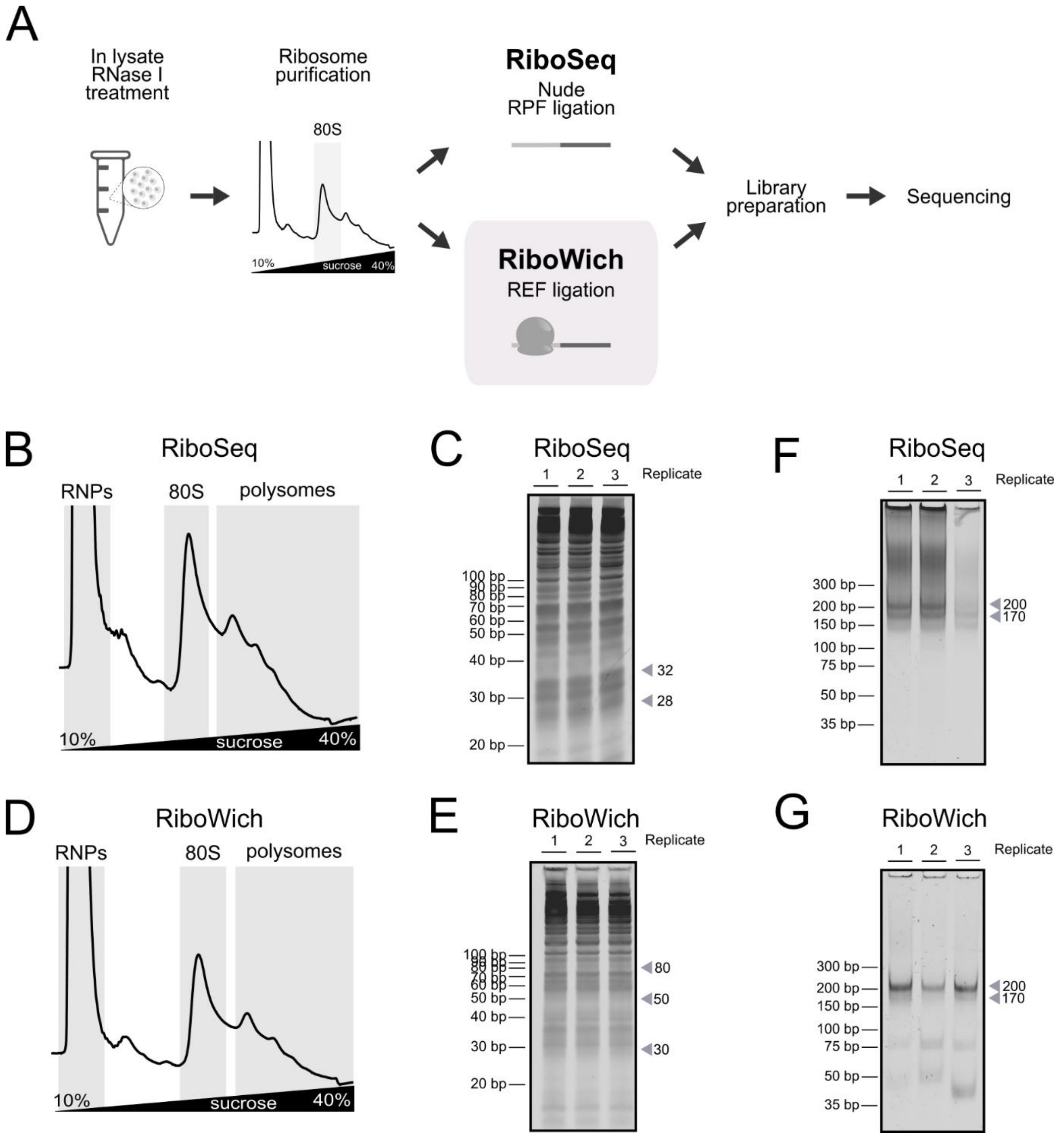


**Supplementary Figure 1**. (**A**) Experimental workflow for in parallel RiboSeq and RiboWich. (**B**) Representative polysome profiles from HEK293T cellular lysates treated with RNase I and used for RiboSeq library preparation. (**C**) Gel electrophoresis for size selection and purification of the RPFs (~30 nt) obtained from three replicates, used for RiboSeq library preparation. (**D**) Representative polysome profiles from HEK293T cellular lysates treated with RNase I and used for RiboWich library preparation. (**E**) Gel electrophoresis for purification of the RiboWich ligation product (~80 nt) for the three replicates. (**F-G**) Gel electrophoresis for the purification of the final libraries (~200 bp) for the three replicates of RiboSeq (top), and RiboWich (bottom).

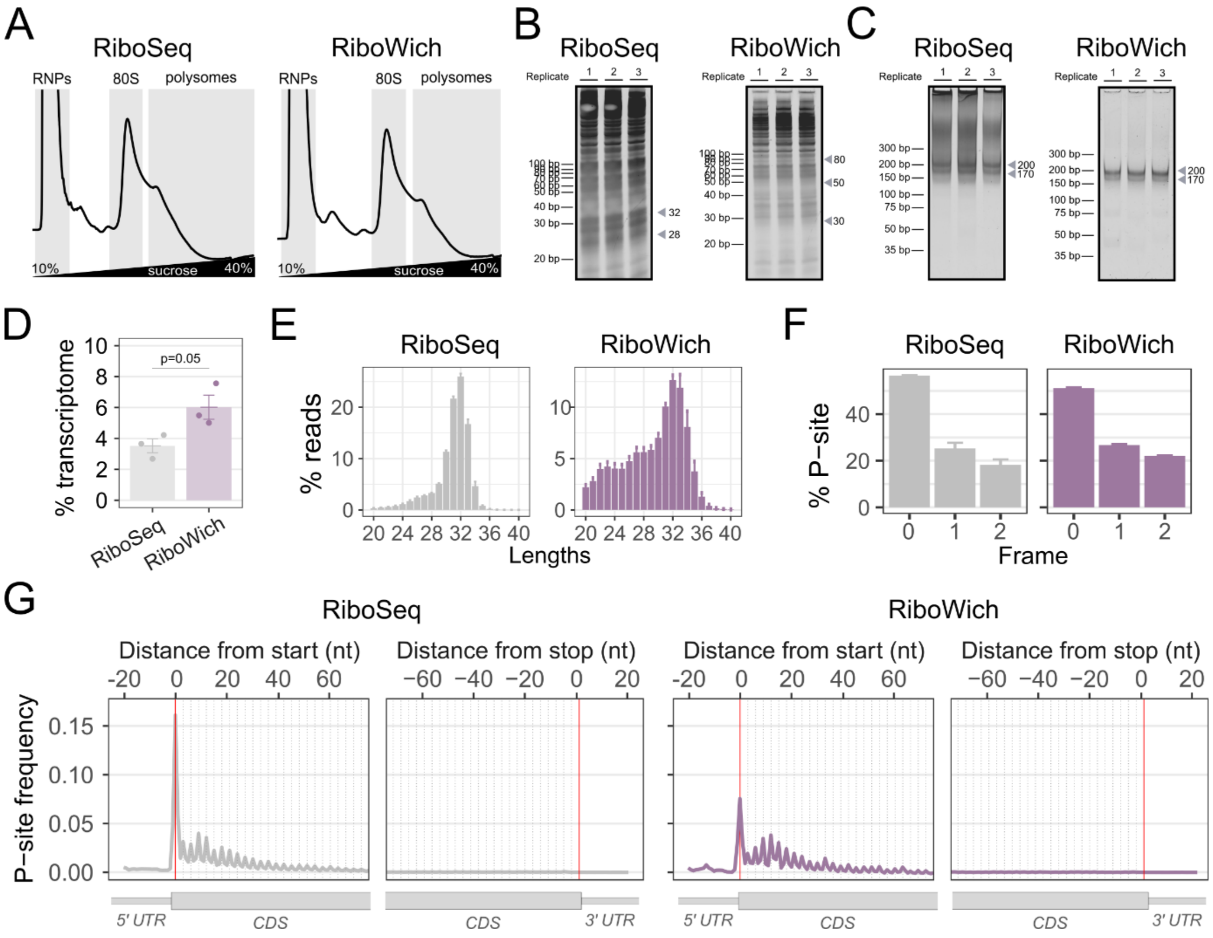


**Supplementary Figure 2**. **Comparison between RiboSeq and RiboWich performed in HEK293T treated with harringtonine**. (**A**) Representative polysome profiles from HEK293T cellular lysates treated with harringtonine and RNase I and used for RiboSeq (left) and RiboWich (right). (**B**) Gel electrophoresis for size selection and purification of RPFs (~30 nt) in RiboSeq (left) and purification of the ligation products (~80 nt) in RiboWich (right) for the three replicates. (**C**) Gel electrophoresis for the purification of the final library (~200 bp) obtained with the RiboSeq (left) and RiboWich (right) for the three replicates. (**D**) Percentages of reads aligning to the transcriptome for RiboSeq and RiboWich. Results represent the mean ± SEM of n=3 independent technical replicates. Two-sided T-test: P-value = 0.05. (**E**) Percentage distribution of read lengths in RiboSeq (left) and RiboWich (right). Results represent the mean ± SEM of n=3 independent technical replicates. (**F**) Percentage of P-sites mapping to the three reading frames for RiboSeq (left) and RiboWich (right). Results represent the mean ± SEM of n=3 independent technical replicates. (**G**) Meta-profiles showing the frequency of P-sites mapping at the beginning and the end of the coding sequence in RiboSeq (left) and RiboWich (right). Results represent the mean ± SEM of n=3 independent technical replicates.

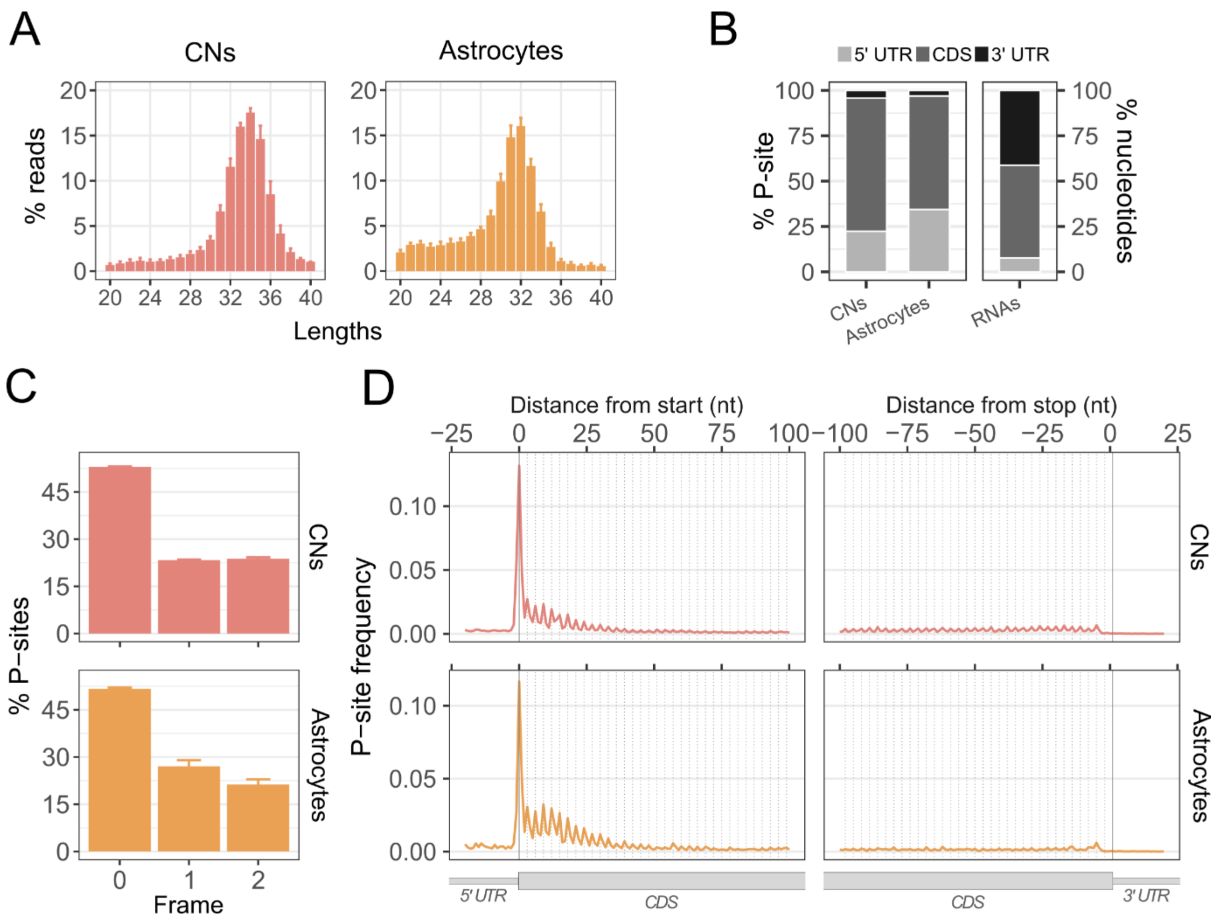


**Supplementary Figure 3**. **RiboWich in CNs and astrocytes samples treated with harringtonine**. (**A**) Percentage distribution of read lengths for CNs (left) and astrocytes (right). Results represent the mean ± SEM of n=2 (CNs) or n=3 (astrocytes) independent technical replicates. (**B**) Percentages of P-sites mapping on the 5' UTR, CDS, and 3' UTR for CNs (left) and astrocytes (middle). Percentages of region lengths in mRNA sequences are reported as reference (right). Results represent the mean of n=2 (CNs) or n=3 (astrocytes) independent technical replicates. (**C**) Percentage of P-sites mapping to the three reading frames for CNs (top) and astrocytes (bottom). Results represent the mean ± SEM of n=2 (CNs) or n=3 (astrocytes) independent technical replicates. (**D**) Meta-profiles showing the frequency of P-sites mapping at the beginning and the end of the coding sequence in CNs (top) and astrocytes (bottom). Results represent the mean ± SEM of n=2 (CNs) or n=3 (astrocytes) independent technical replicates.

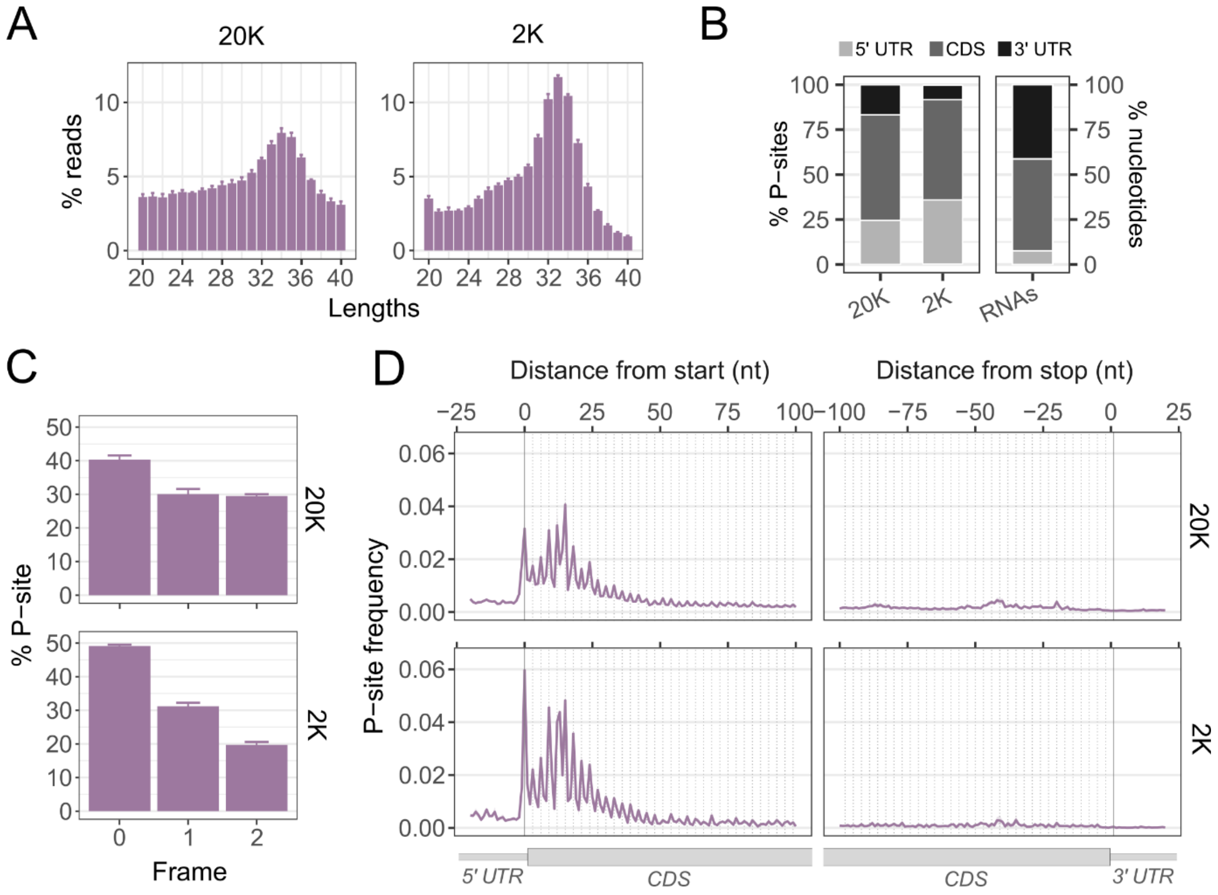


**Supplementary Figure 4**. **Low-input RiboWich in samples treated with harringtonine: comparison between 20,000 (20K) and 2,000 (2K) HEK293T**. (**A**) Percentage distribution of read lengths for 20K (left) and 2K (right). Results represent the mean ± SEM of n=6 (20K) or n=18 (2K) independent technical replicates. (**B**) Percentages of P-sites mapping on the 5' UTR, CDS, and 3' UTR for 20K (left) and 2K (middle). Percentages of region lengths in mRNA sequences are reported as reference (right). Results represent the mean of n=6 (20K) or n=18 (2K) independent technical replicates. (**C**) Percentage of P-sites mapping to the three reading frames for 20K (top) and 2K (bottom). Results represent the mean ± SEM of n=6 (20K) or n=18 (2K) independent technical replicates. (**D**) Meta-profiles showing the frequency of P-sites mapping at the beginning and at the end of the coding sequence in 20K (top) and 2K (bottom). Results represent the mean ± SEM of n=6 (20K) or n=18 (2K) independent technical replicates.

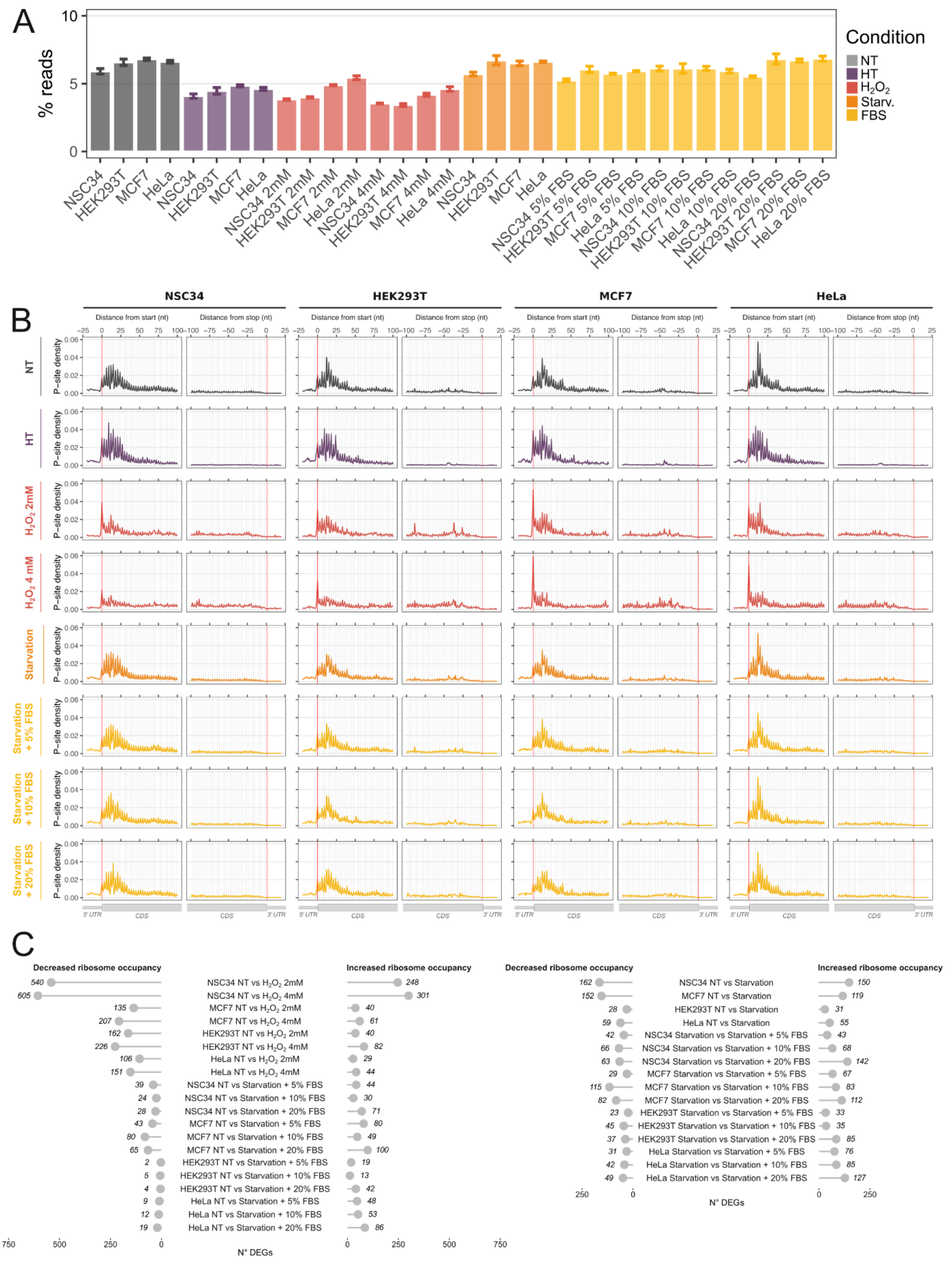


**Supplementary Figure 5. High-throughput RiboWich with 96 samples**. (**A**) Percentage of reads mapping to the transcriptome. Results represent the mean of n=3 technical replicates for each condition and each cell line. (**B**) Meta-profiles of P-site distribution at the start and end of coding sequences in the different conditions and cell lines. Results represent the mean ± SEM of $n = 3$ independent technical replicates for each condition and cell line. (**C**) Number of genes showing significant changes in ribosome occupancy for each condition and each cell line (|fold change| > 0.5, adjusted $p < 0.05$).
